# Quantifying uncertainty in RNA velocity

**DOI:** 10.1101/2024.05.14.594102

**Authors:** Huizi Zhang, Natalia Bochkina, Sara Wade

## Abstract

The concept of RNA velocity has made it possible to extract dynamic information from single-cell RNA sequencing data snapshots, attracting considerable attention and inspiring various extensions. Nonetheless, existing approaches lack uncertainty quantification and many adopt unrealistic assumptions or employ complex black-box models that are difficult to interpret. In this paper, we present a Bayesian hierarchical model to estimate RNA velocity, which leverages a time-dependent transcription rate and non-trivial initial conditions, allowing for well-calibrated uncertainty quantification. The proposed method is validated in a comprehensive simulation study that covers various scenarios, and benchmarked against a widely embraced and commonly recognized approach for RNA velocity on single-cell RNA sequencing data from mouse embryonic stem cells. We demonstrate that our model surpasses this widely used, state-of-the-art method, offering enhanced interpretation of cell velocity and cell orders. Additionally, it supports the estimation of a unified gene-shared latent time, providing a valuable resource for downstream analysis.

## 1 Introduction

Single-cell RNA sequencing (scRNA-seq) is a powerful tool that enables measurements of transcriptome profiles for individual cells across thousands of genes. It has been used to provide more comprehensive understanding of individual cells, such as identifying subpopulations, understanding mechanisms underlying cellular differentiation, studying cancers and other biological processes (Tang et al., 2019). However, since the cells are killed when measurements are taken, the resulting scRNA-seq data only provides a static snapshot of cellular states. To study dynamic information of the cells, La Manno et al. (2018) proposed the notion of RNA velocity that can be derived from a per-gene *steady-state* model, relying on the amount of unspliced mRNAs, denoted by *u*, compared to spliced mRNAs, denoted by *s*. Both *s* and *u* can be identified from the total mRNAs, by leveraging the fact that *u* is characterized with introns, and after removing introns, the remaining adjacent exons are connected to produce *s*.

For every gene, RNA velocity of a cell is defined as the first time derivative of the spliced mRNA, indicating the direction of the change of spliced counts. Positive velocity implies the gene is being induced (induction state), resulting in more unspliced mRNA than expected in steady state, whereas negative velocity (repression state) suggests the gene is being down-regulated with less unspliced mRNA than expected. Hence, combining RNA velocities across all genes, it is possible to predict the future states of cells.

The steady-state model, implemented in *velocyto*, assumes that the cells have reached equilibrium in the induction phase, which may not necessarily hold true in practice. The *dynamic model* available in *scVelo* (Bergen et al., 2020) relaxes such assumptions, utilizing the Expectation-Maximization (EM) algorithm to infer parameters. Furthermore, it provides an estimate for latent time, denoted by *t*, per cell and per gene, describing the position of the cell in the biological process being studied.

The two models have become the crucial basis of numerous extensions that have emerged recently. Both models are underpinned by the assumption that the gene is first transcribed to unspliced mRNA with a state-dependent transcription rate *α*, then unspliced mRNA is spliced out with a constant splicing rate *β* to produce spliced mRNA, followed by a constant degradation rate *γ* capturing the downregulation of spliced mRNA, where all the rates are gene-specific.

Despite of the popularity of the two methods, they are known to suffer from several limitations. First, neither quantifies the uncertainty in the velocity estimates or latent times, and therefore it is difficult to understand how reliable the velocity estimates are for down-stream analysis and interpretation. Secondly, the assumptions of a constant transcription rate within each state and constant splicing and degradation rates may not be adequate for complex real data, as suggested by Bergen et al. (2021). Possible solutions include manual removal of MUltiple Rate Kinetics (MURK) genes that violate model assumptions (Barile et al., 2021). Additionally, the relationship between *u* and *s* may not be correctly captured by the dynamic model due to the complexity of the biological process (Riba et al., 2022).

In this work, we develop a Bayesian hierarchical model to estimate RNA velocity along with kinetic rate parameters to address these problems. Bayesian methods have been proven to be instrumental to capture the variability inherent in the single-cell data (Huang and Sanguinetti, 2021). Our primary innovation lies in a full Bayesian treatment for sound uncertainty quantification, along with the introduction of a time-dependent transcription rate and the flexibility of allowing for a non-zero initial condition, generalizing the dynamic model. Moreover, we introduce the concept of *consensus velocity*, an algorithm which can better account for the uncertainty in the state and can be easily adapted for Markov chain Monte Carlo (MCMC) methods. We compare our method *ConsensusVelo* applied to the real scRNA-seq data from embryonic stem cells (Riba et al., 2022) with scVelo and show that our approach outperforms scVelo in capturing the relationship between spliced and unspliced mRNA, and also provide a comparison with a deep learning based state-of-the-art method DeepCycle (Riba et al., 2022). Finally, we propose a fast Bayesian model for estimating a gene-shared latent time using posterior summaries from ConsensusVelo, while also capturing the associated uncertainty. Due to high variability between cells, quantifying the uncertainty in the temporal ordering of cells is of great importance; indeed, (Campbell and Yau, 2016) demonstrate how using a point estimate may result in increased false discovery rates in downstream differential analysis.

## 2 Review of Dynamic Modelling of RNA Velocity

We begin our review with the dynamic model of Bergen et al. (2020) in Section 2.1, which forms the basis of our proposed model, followed by an overview of extensions in Section 2.2.

### 2.1 Dynamic Model with Constant Transcription Rate

RNA velocity *v*(*t*) is defined as the rate of change of spliced RNA *s*(*t*): *v*(*t*) = *ds*(*t*)*/dt*. The most simple differential equations that describe the rates of change over time of spliced and unspliced RNA, *s*(*t*) and *u*(*t*), for every gene are given by

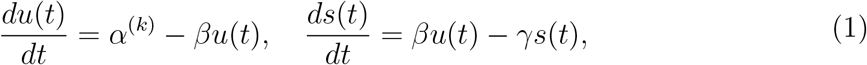

with *k* = 1 for induction and *k* = 2 for repression. Note that the transcription rates *α*^(*k*)^, as well as splicing and degradation rates (*β* and *γ*) here are assumed to be constant with time. The full solutions to the differential equations are analytically available in Bergen et al. (2020), with the introduction of a starting time point for each state 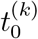, and the time a cell *c* has spent in its associated state *k*_*c*_, defined as 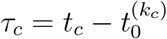.

Let *x*(*t*) = (*u*(*t*), *s*(*t*)) and 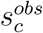 and 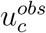 denote the (processed) observed counts for cell *c* in a particular gene. The residuals defined as signed Euclidean distance 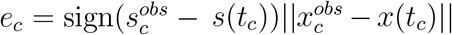 are assumed to follow a normal distribution N(0, *σ*^2^) with gene-specific variance *σ*^2^. Note that in order to improve data quality and use the normality assumption, the observed counts are first processed (as described in Bergen et al. (2020)). The EM algorithm consists of allocating a latent time to each cell that minimizes the distance to *x*(*t*) in the E-step, and then maximizing the log-likelihood with respect to rates and 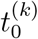 in the M-step.

### 2.2 Extensions

Numerous other methods have also been proposed to address the limitations of scVelo and enhance the reliability of RNA velocity modeling. Qiu et al. (2022) obtain more accurate velocity inference through RNA metabolic labeling, enabling the analysis of time-resolved single-cell RNA-seq data and the construction of vector fields for predicting cell fates.

Recently, the potential to understand the dynamics without further calibrated experiments has garnered significant attention. Gorin et al. (2022) point out that both the steady-state model and the dynamic model are not rich enough to account for bursty transcription, and argue for alternative approaches based on chemical master equations. UniTVelo (Gao et al., 2022) generalizes the relationship between *s* and latent time *t* by a radial basis function. There are approaches that avoid using the dynamic model entirely, e.g. Velo-Predictor (Wang and Zheng, 2021), an ensemble machine learning pipeline, which treats RNA velocity as a classification problem. To avoid a separate pre-processing step, Qin et al. (2022) construct an end-to-end probabilistic generative model Pyro-Velocity in a Bayesian framework based on raw counts.

Additionally, recent research has increasingly harnessed the power of neural networks. Several approaches, such as DeepVelo (Cui et al., 2022), CellDancer (Li et al., 2023) and VeloVAE (Gu et al., 2022), generalize the rate parameters to be cell-specific. VeloAE (Qiao and Huang, 2021) finds a low-dimensional representation for *u, s, v* and filter noise with an autoencoder. LatentVelo (Farrell et al., 2022) aims to capture the causal relationship between the unspliced and spliced RNA, going beyond the linear differential equations. VeloVI (Gayoso et al., 2022) is a Bayesian deep generative model assuming a time-dependent transcription rate. Although VeloVI shares a similar functional form for the transcription rate *α* with our proposed model during the induction phase, it keeps *α* constant during repression. On the contrary, our model specifies a smooth rate parameter that applies to both induction and repression.

With regard to estimating the uncertainty of the parameter estimates and of the RNA velocity, all the above approaches provide their point estimates. LatentVelo, VeloVAE, VeloVI and Pyro-Velocity output a variational posterior distribution which in principle can estimate the uncertainty however this approach is known to significantly underestimate it.

## 3 Model

### 3.1 Dynamic Model with Time-dependent Transcription Rate

The assumption of constant transcription rate can lead to inflated curvature and poor fit (Figure 1 top) and widespread changes in kinetic rates over time have been observed during cell cycle (Zopf et al., 2013; Battich et al., 2020), thus encouraging a more flexible model to understand transcription mechanism.

**Figure 1.**
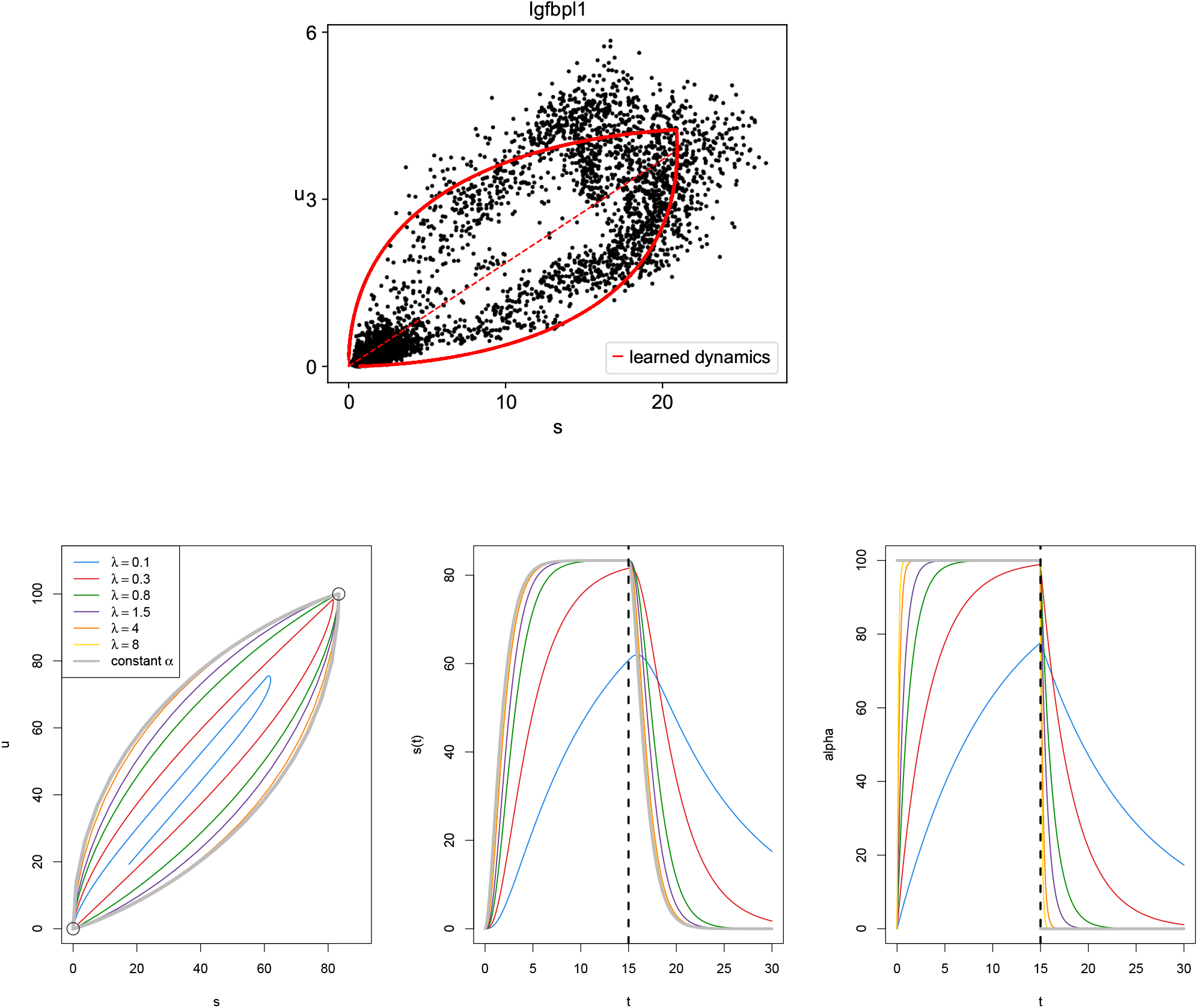
Top: Fitted phase portrait from scVelo for gene *Igfbpl1* in Pax6 data (Manuel et al., 2022). Bottom: Comparison between varying values of *λ*, and the const-*α* model. Bottom-left: phase portrait. Bottom-middle: *s*(*t*) against latent time *t*. Bottom-right: time-dependent transcription rate *α*(*t*) against *t*. The black circles on the left panel are the steady-state values. The black dashed line on the right two panels is the switching point 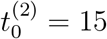.

**Figure 2.**
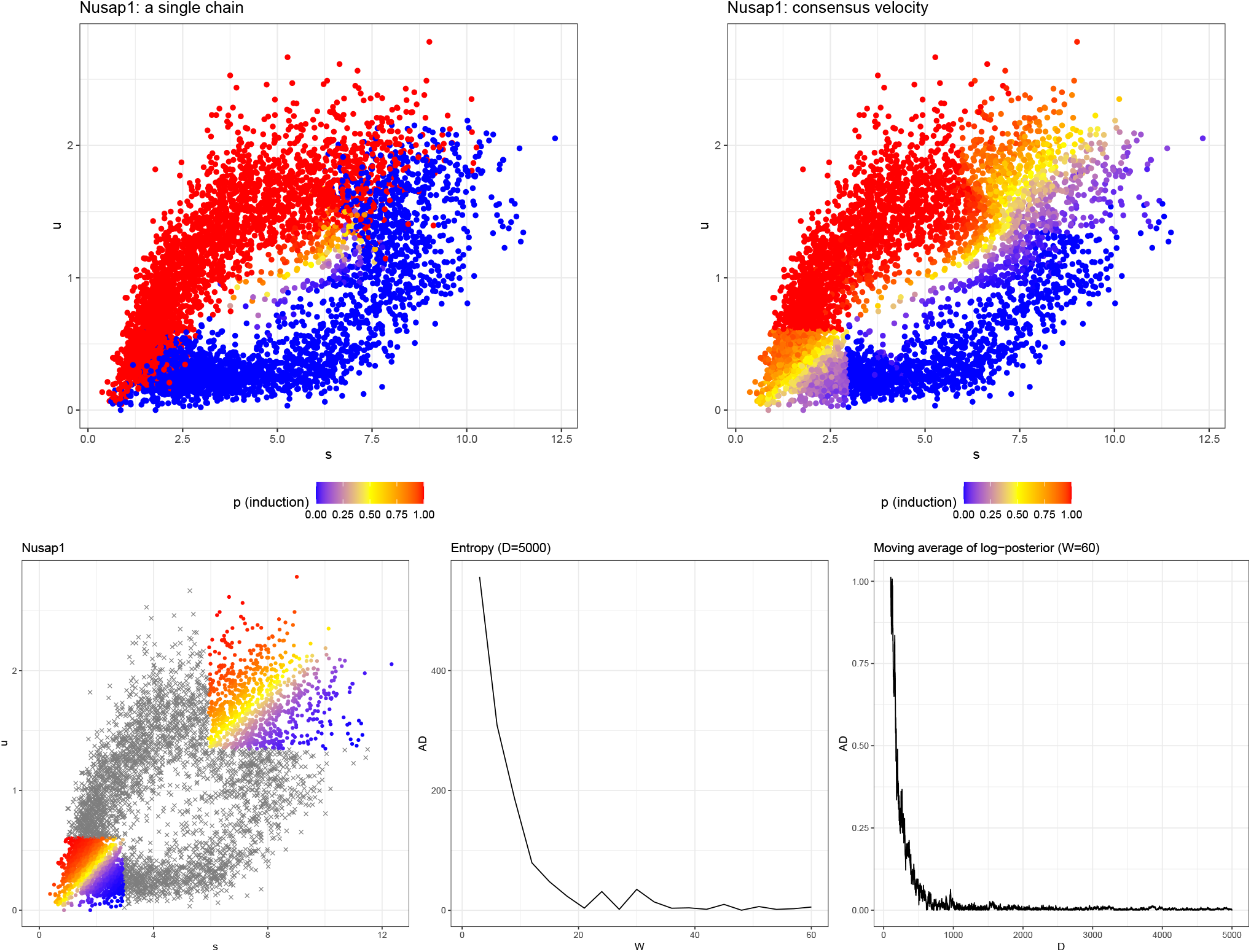
Top: Posterior probability of induction for gene *Nusap1* in data collected from mouse embryonic stem cells (Riba et al., 2022), using a single chain (left) and consensus velocity (right). Bottom-left: Empirical probabilities of belonging to induction used for multiple initializations of uncertain cells in ConsensusVelo for gene *Nusap1*. Bottom-middle: Absolute difference in entropy for *W* = (3, 6, …, 60), given *D* = 5000. Bottom-right: Absolute difference in moving average of log-posterior across all 60 chains for *D* = (101, 102, …, 5000).

Motivated by Bergen et al. (2021), we extend the dynamic model (Eq (1)) by allowing *α* to be a smooth function over time

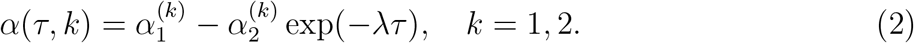

By substituting Eq (2) into Eq (1), the closed form solutions for our proposed model are:

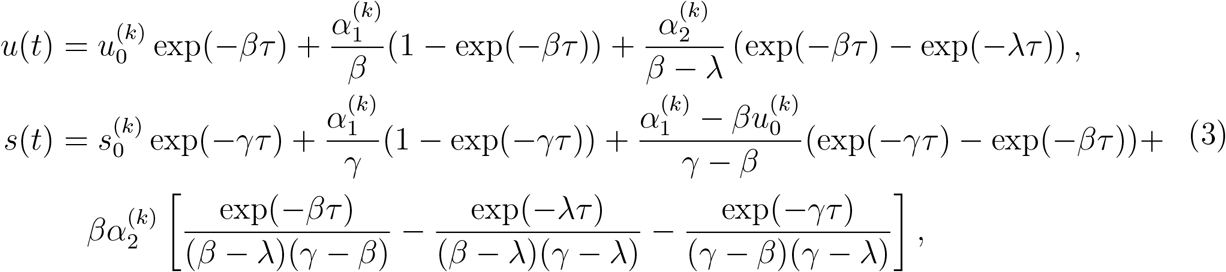

where 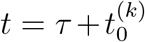 and *β≠ γ≠ λ*. When either two of *β, γ, λ* are equal, it is also possible to obtain an analytic solution. To ensure that *u*(*t*) and *s*(*t*) are continuous, the initial values 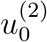 and 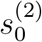 satisfy

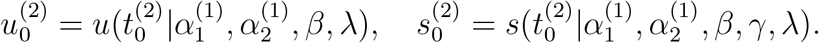

When each state lasts for a sufficiently long time (*τ → ∞*), the abundances reach the steady state

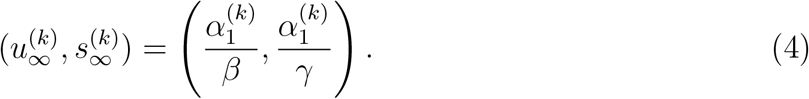

Note that if 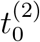 is large enough, the initial conditions 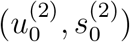 will be approximately equal to the induction steady-state values.

*u*(*t*) and *s*(*t*) under the proposed model are plotted in Figure 1 (bottom) under various values of *λ*. As *λ* increases, *u*(*t*) and *s*(*t*) closely resemble those in the dynamic model (referred to as the const-*α* model afterwards), with *α*(*t*) becoming nearly constant 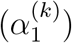. On the other hand, for small *λ* = 0.1, the phase portrait of *u* versus *s* approximates a straight line with minimal curvature. As *λ* decreases, the upper right steady state is less likely to be reached, similar to the effect of a small 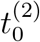 (known as “early switching” in Bergen et al. (2020)).

#### 3.1.1 Assumptions

We list our assumptions for the proposed model (for a single gene):

1. We assume that 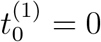. Consequently, *k*_*c*_ = 1 if 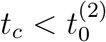, and *k*_*c*_ = 2 otherwise.
2. The transcription rate *α*(*t*) is assumed to be continuous. This holds if it is continuous at 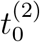, which represents the end point of the induction state and the starting time of the repression state; thus, we require 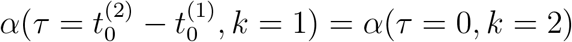, i.e.,

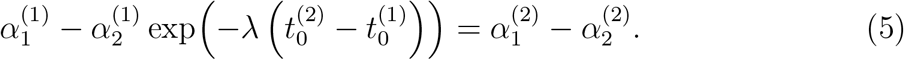
3. Unlike the const-*α* model, the initial conditions 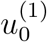 and 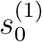 are assumed unknown. In such cases, in order to have a closed path, i.e., the repression steady state matches the initial condition, it is required that 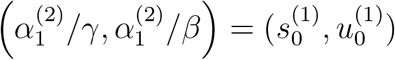,i.e.,

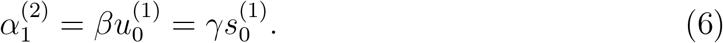

This constraint implies that the points (0,0), initial condition (equivalently the repression steady state) and the induction steady state 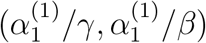 are on the same line. Since the point 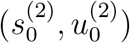 approximates the induction steady state when the induction is sufficiently long, one would expect the origin and the two steady states of the observed phase portrait to lie on the same line. In addition, for a non-zero initial condition, it is necessary that 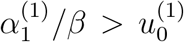 to establish that the induction steady state is above the repression steady state in the phase portrait, which is automatically satisfied when 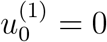.
4. At *t* = 0, assume the first time derivative of the unspliced count is zero: 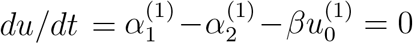. Combining this with the assumption of continuity of *α* (Eq (5)) yields

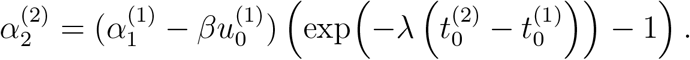

These assumptions greatly reduce the number of unknown parameters. Specifically, we only need to estimate one initial condition (Eq (6)), and only 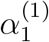 instead of 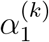 and 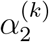(*k* = 1, 2), as well as *γ, λ*, 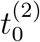, *τ*_*c*_ and *k*_*c*_.

### 3.2 Likelihood and Identifiability

We assume the following model for the observed unspliced count 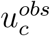 and spliced count 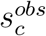 (for a single gene) pre-processed in scVelo (Bergen et al., 2020):

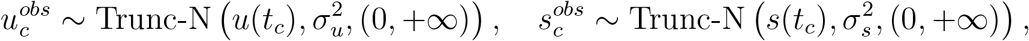

where *u*(*t*_*c*_) and *s*(*t*_*c*_) are given by Eq (3) and Trunc-N (*μ, σ*^2^, *A*) denotes the truncated normal distribution with mean *μ*, variance *σ*^2^ and truncation region *A*. The truncation is applied to account for the positive nature of the (processed) counts.

Similar to the const-*α* model, the parameters are identifiable only up to a constant (scale invariance). For instance, if 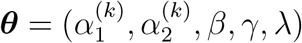 is one solution, then

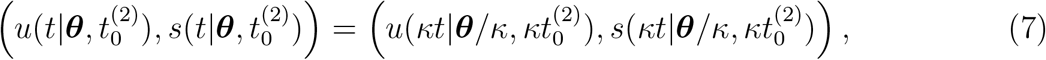

for any *κ >* 0. For identifiability, we choose to parametrize the model such that the rates and time-relevant quantities are measured relative to *β*.

Conditional on the other parameters, large *τ*_*c*_ can only be weakly identified from the likelihood, since the cell approaches the steady state with approximately constant mean values *u*(*t*_*c*_) and *s*(*t*_*c*_) (Figure 1 bottom). The likelihood then remains almost constant for any sufficiently large *τ*_*c*_ (see discussion in terms of the Fisher information of *t*_*c*_ in Web Appendix A). The same issue applies for large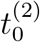. Lastly, such weak identifiability is also present for large *λ* because the values *u*(*t*) and *s*(*t*) will approximate those from the const-*α* model.

### 3.3 Empirical Parameter Estimates

Now we discuss how to obtain empirical estimates for the model parameters, some of which coincide with those of Bergen et al. (2020).

First we identify the sets of cells in the steady states: ON for the end of the induction, and OFF for the end of repression, assuming that cells in the upper right corner of the phase portrait are in the induction steady state (Bergen et al., 2020) (see Figure 3 in Web Appendix B for an example gene). Note that this assumption may not always be true, particularly in the case of a small 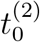 or *λ*. Then, given that the velocity for the steady state is approximately zero, the estimated degradation rate is 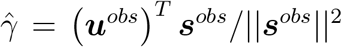 (recall that each parameter is measured relative to *β*), where ***u***^*obs*^ and ***s***^*obs*^ denote observations in the steady states. The rate 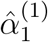 is calculated based on the steady-state values (Eq (4)), specifically set to the maximum value of ***u***^*obs*^ (or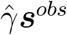). The initial conditions 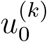 and 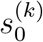 are estimated in a similar way using steady-state cells (see Web Appendix C). As for the standard deviations, the empirical values are derived as follows:

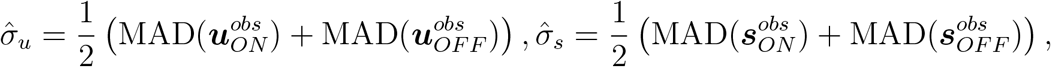

where MAD denotes the median absolute deviation which is more robust than standard deviation. The state is estimated as 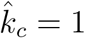 if 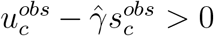 and 2 otherwise. Finally, for *τ*_*c*_, we invert *u*(*t*) in the const-*α* model (replaced with its observed value), to obtain

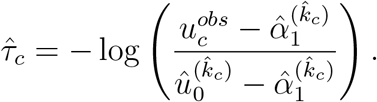

**Figure 3.**
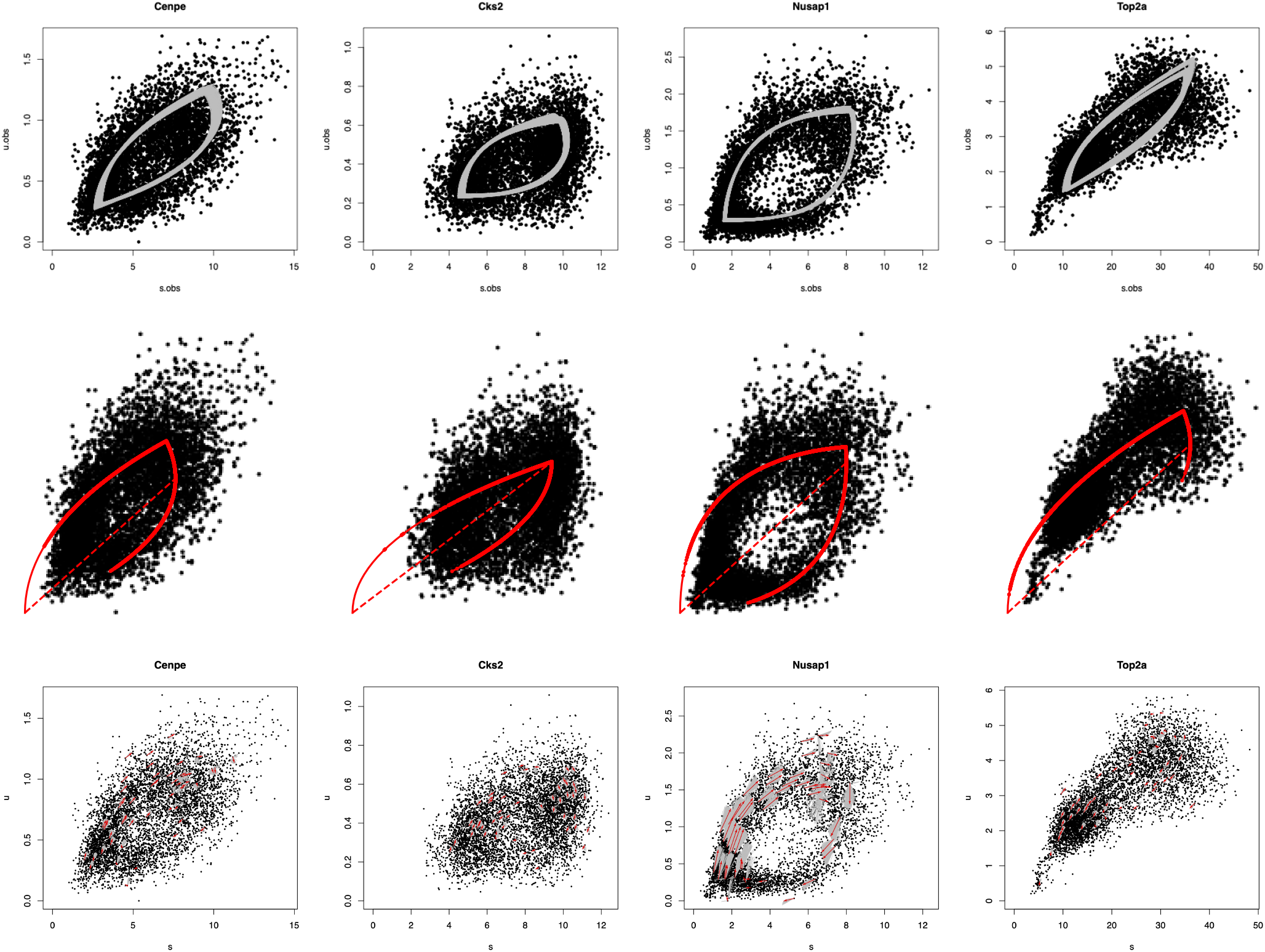
Fitted phase portrait for example genes with uncertainty (200 MCMC samples in grey) from ConsensusVelo (top) and scVelo (middle) (Riba et al., 2022; Bergen et al., 2020). The bottom row shows predicted velocity for example cells, with uncertainty (200 MCMC samples in grey; red arrow denotes posterior mean).

**Figure 4.**
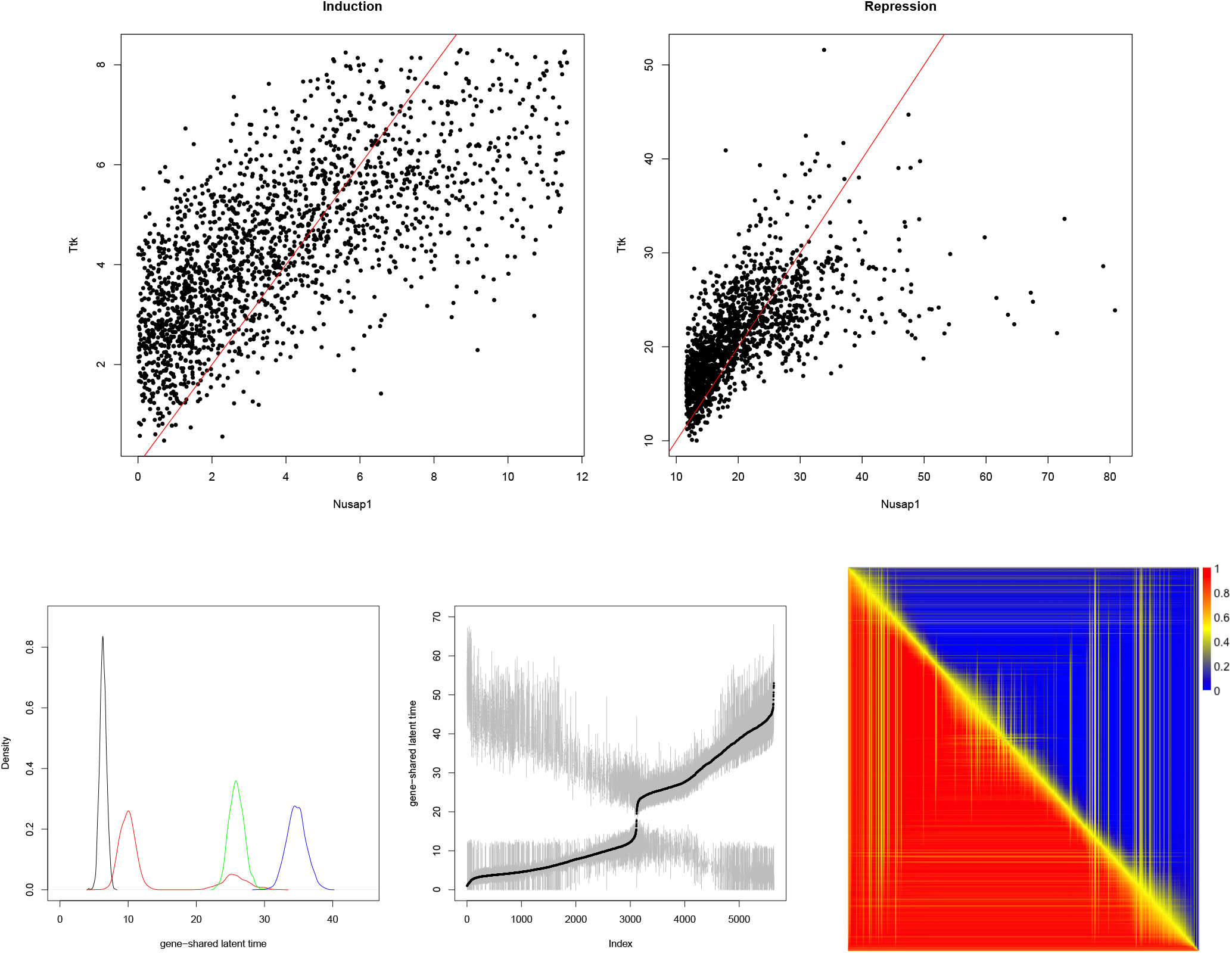
Top: Comparison of gene-specific latent time between two genes, conditional on the state. The red line denotes *y* = *x*. Bottom-left: Posterior densities of gene-shared latent time for selected cells. Bottom-middle: 95% HPD CI (grey) and posterior median (black) of shared latent time. Cells are ordered by increasing posterior median of 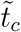. Bottom-right: Posterior probability of one cell before another; entry (*i, j*) is the posterior probability of the cell in row *i* before the cell in column *j*. Cells are ordered by decreasing posterior median of 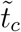 from top to bottom, and left to right.

It is possible that the term inside the log may be negative or greater than 1; in such cases, *−* log(*ϵ*) (or *−* log(1 *− ϵ*)) is used as the value for 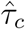, for small positive *ϵ*. Given 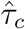, we estimate 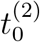 as 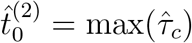 among cells with 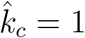 only.

### 3.4 Prior Distributions

We use a Bayesian approach to infer parameter values. The unknown local time *τ* is modeled exchangeably in each state:

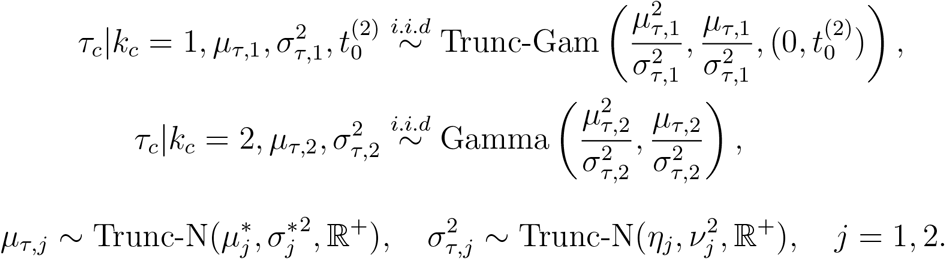

Since only two observations contribute to the likelihood of each *τ*_*c*_, and large *τ*_*c*_ are only weakly identifiable (Section 3.2), it is crucial to use an exchangeable hierarchical prior for *τ*_*c*_ which allows to borrow information within each state; see also Web Appendix A. The remaining parameters have fairly standard prior distributions, with the hyperparameters fixed to their empirical values, and *ξ* = 1 (see Web Appendix C for model details):

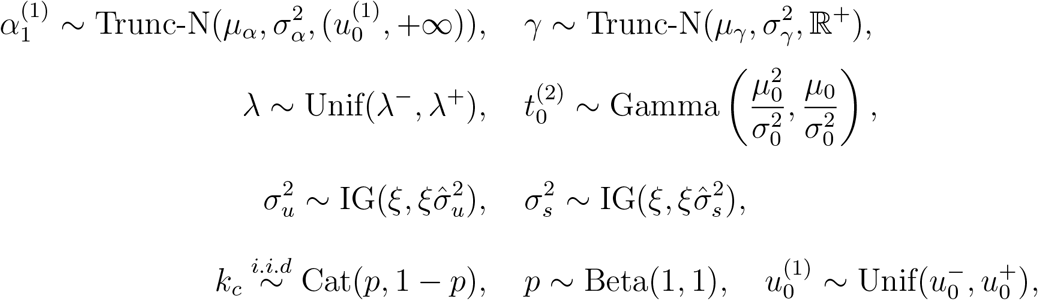

where *p* is the hyperparameter denoting the probability of being in the induction phase. A graphical model representation is shown in Figure 5 in Web Appendix D.

**Figure 5.**
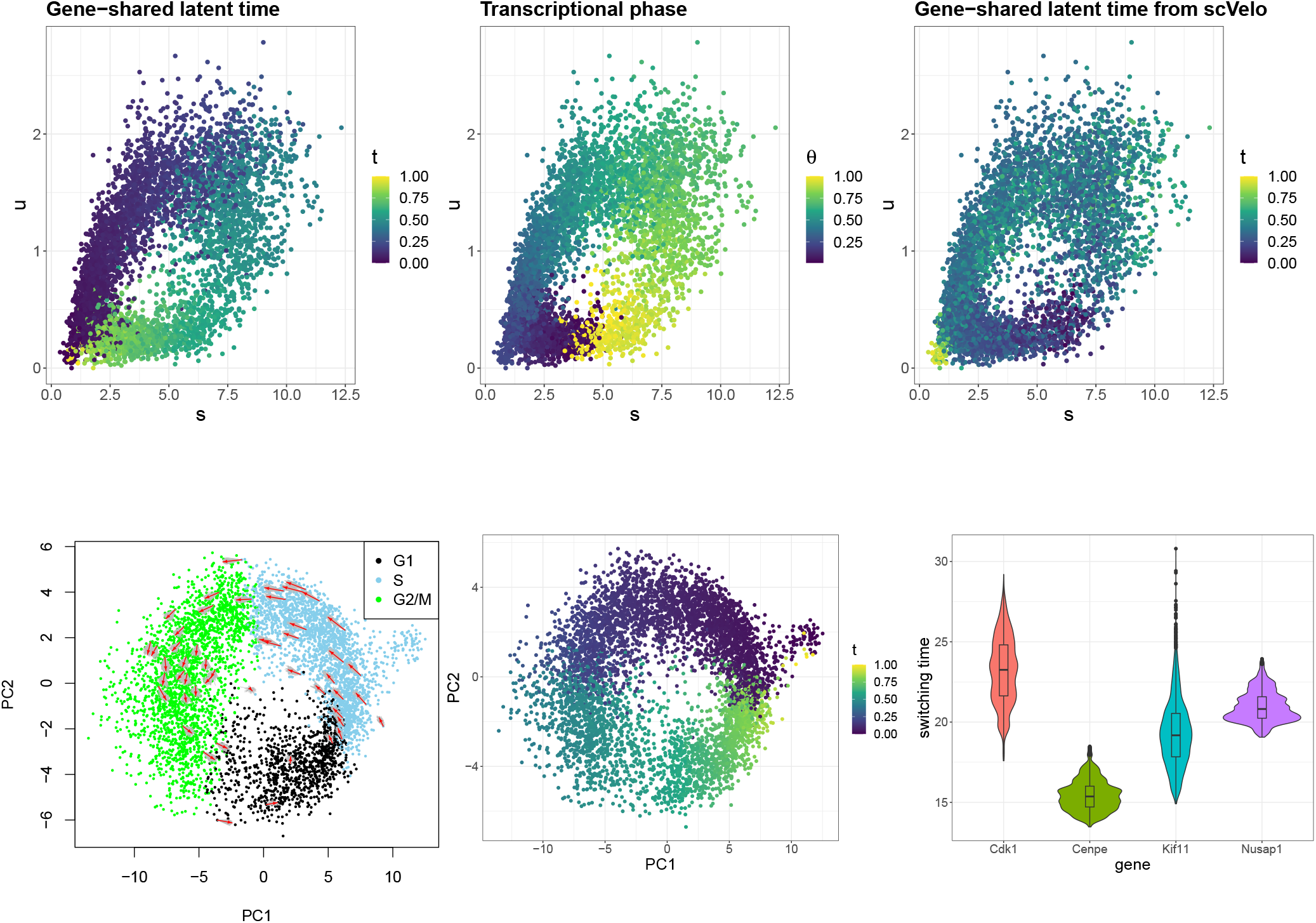
Top: Comparison of gene-shared latent time from ConsensusVelo (left) with transcriptional phase *θ* from DeepCycle (middle) and latent time from scVelo (right) on gene *Nusap1*. Bottom-left: Predicted velocity with uncertainty (results are based on 200 MCMC samples) for example cells on the PCA space, after rescaling. The cells are colored by cell cycle phases. Bottom-middle: Posterior median of gene-shared latent time 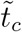on the PCA space. Bottom-right: Comparison of switching time across different genes. Posterior densities of adjusted 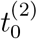 are plotted for examples genes.

## 4 Posterior Inference

We develop a Metroplis-within-Gibbs algorithm to draw posterior samples. For *k*_*c*_ and *p*, the full conditional distributions can be sampled from directly, while for all other parameters, adaptive Metropolis-Hastings (AMH) is implemented for each parameter individually. To target an average acceptance probability of 0.44 that is optimal for univariate variables (Roberts and Rosenthal, 2009), the AMH scheme described in Algorithm 5 from Griffin and Stephens (2013) is applied. The algorithm is given in Web Appendix D.

### 4.1 MCMC: Challenges and Proposed Solutions

It is worth noting that the uncertainty in *k*_*c*_ and *τ*_*c*_ for cells located near the two corners (upper-right and lower-left area in the phase portrait) may be underestimated (Figure 2 top-left). For these cells (referred to as uncertain cells hereafter), their associated states are expected to have more uncertainty than the other cells, as they may belong to either of the two states. For instance, upper-right uncertain cells may be in *k* = 1 with *τ* close to 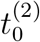, or in *k* = 2 with *τ* close to zero. However, in the Gibbs sampling step for *k* that is conditioned on *τ*, proposing a cell at the end of the induction with *k*_*c*_ = 1 and 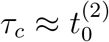 to *k*_*c*_ = 2 with the same *τ*_*c*_ that could be substantially larger than zero, is incorrect. Such a proposal is highly unlikely to be accepted, resulting in sampled states trapped in one single mode.

Secondly, due to the high dimensionality (cell-wise parameters *τ* and *k*), the Gibbs sampling can face the dilemma of getting stuck in local modes, and running for a long time might yield minimal improvement. Moreover, because of the weak identifiability issue discussed in Section 3.2, the chains are very likely to converge to different places, whereas the likelihood (or posterior) may already stabilize. Therefore, standard diagnostic methods to check convergence for MCMC chains are no longer appropriate.

To mitigate these limitations, we propose the idea of consensus velocity, motivated by consensus clustering (Coleman et al., 2022) which addresses similar issues in clustering. We refer to our model and algorithm together as *ConsensusVelo*, which instead of running a few chains for a long time, considers a large number of chains (*W*) with different initial *k* for the uncertain cells (see Figure 2 bottom-left and more details in Web Appendix B) and runs each chain for a suitable number of iterations *D*. Usually *D* is small in the case of high dimensionality. This approach allows for parallelization, yielding less computational cost. The samples from the last part of the *W* chains are combined for posterior inference.

The critical step of consensus velocity is choosing the tuning parameters *W* and *D*. Let (*w*_1_, …, *w*_*J*_) be candidate values of *W* in increasing order. The number of chains *W* needs to be large enough to account for the uncertainty in *k*. For a given number of iterations *d*, we compute the vector ***p*** = (*p*_1_, …, *p*_*C*_) representing the probability of being in the induction state for each cell *c ∈* [1, …, *C*] and entropy *E*, based on the samples for *k* at the *d*-th iteration from the *w*_*j*_ chains, as follows:

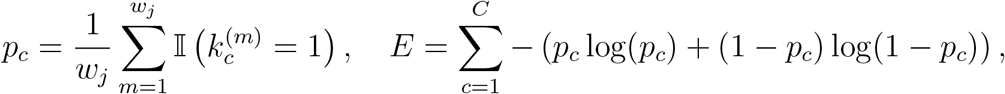

where 𝕀(*·*) denotes the indicator function, and 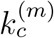 is the sample for *k*_*c*_ from the *m*-th chain (*m* = 1, …, *w*_*j*_). The entropy *E* describes the total uncertainty in states for all cells, and we compute the absolute difference (AD) in *E* from *w*_*j*_ chains to that from *w*_*j−*1_ chains. In the beginning, adding chains with different initial *k* introduces more uncertainty in cell states and hence AD is expected to be large, but as more chains are included, we have fewer gains. Thus, the plot of these absolute differences against *w*_*j*_ is expected to have an elbow shape, and a suitable *W* is chosen at which the curve plateaus. As for *D*, denote the candidate values by (*d*_1_, …, *d*_*I*_). For a fixed number of chains *w*, we monitor the moving average of the log-likelihood or log-posterior across all *d*_*i*_ iterations and *w* chains. The absolute difference in the moving average from *d*_*i*_ to *d*_*i−*1_ iterations is used to determine *D*. An example of choosing *W* and *D* is provided in Figure 2 (bottom-middle and bottom-right).

Lastly, initialization of *τ*_*c*_ can be substantially influential, and inaccurate initial values can lead to completely erroneous state estimates (see Section 5), resulting in explosively large variance estimates. To address this, we conduct a *preparation step* where *k* is fixed to some sensible empirical estimates, and run the algorithm to obtain samples for all the other parameters, especially *τ*_*c*_. Then the posterior means of these samples will be used as the initial values to run the complete algorithm.

### 4.2 Posterior Predictive Checks

Posterior predictive checks are essential in Bayesian inference to check whether the model, including priors, is a good fit to the observed data. Typically, replicated data is generated given posterior draws of all the unknown parameters to approximate the posterior predictive distribution, and discrepancy between the observed and replicated data is measured to assess goodness-of-fit. For a single replicate, we compare generated observations to true observations directly. For multiple replicates, key statistics such as quantiles of the data and quantities of interests (empirical estimate of *γ*) are compared between simulated and observed data. The posterior predictive p-values (ppp) (Gelman et al., 1996) are widely adopted when conducting posterior predictive checks. For a statistic *T*, the p-value is given by

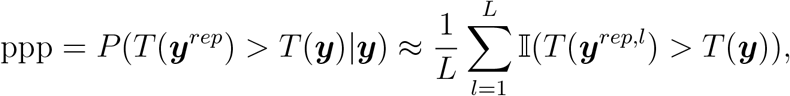

where ***y***^*rep,l*^ denotes one replicate based on a posterior draw.

In the presence of hierarchical models, it is suggested to use a mixed predictive distribution where the parameters are generated from its hyper-priors, yielding less conservative p-values (Marshall and Spiegelhalter, 2003). Besides, for a mixture model as is the case here (two states/components), the p-values should be computed conditional on the component, or only for observations associated with a high posterior probability of being in the component (Lewin et al., 2007). For each state *j*, we compute the p-values as 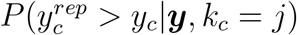 for *s* and *u*, based on cells with *P* (*k*_*c*_ = *j*|***y***) *>* 0.9. The parameter *τ* is generated from its prior given posterior samples of *μ*_*τ,j*_ and 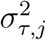, and then the replicate is generated conditional on *τ* and posterior draws for other parameters.

### 4.3 Inference for Gene-shared Latent Time

Given gene-specific latent time, it is natural to consider a unified gene-shared latent time, for which we propose a post-processing step to balance between computational efficiency (i.e. parallel inference for velocity across genes) and appropriate modeling of uncertainty. First, denote 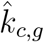 as the state with a larger posterior probability, and 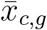 the posterior mean of log *t*_*c,g*_, conditional on 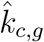. Note that, for every gene *g*, as we measure all parameters relative to *β*_*g*_, the kinetic rate parameters and time-relevant quantities have different scales across genes. The correct gene-specific latent time is *t*_*c,g*_*/β*_*g*_, with the gene-specific kinetic rates corrected by *β*_*g*_ (Eq (7)). The model to infer gene-shared latent time is given by

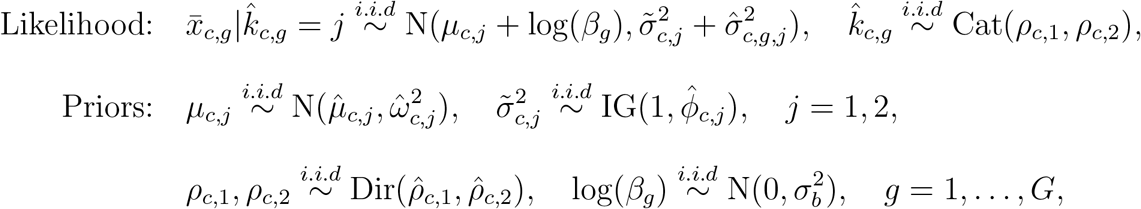

where Cat and IG denote the categorical and inverse-gamma distributions, and 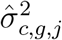 is the posterior variance of log *t*_*c,g*_ conditional on *k*_*c,g*_ = *j*. The variance 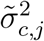 is introduced to model the variability between genes. For identifiability, *β* _1_ is set to 1, meaning every gene is measured relative to the first gene. Note that 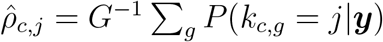 and the posterior mean of *ρ*_*c,j*_ is 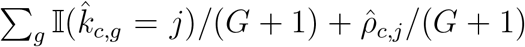, making it more robust to estimating the state *j*. The gene-shared latent time 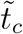 is defined to be exp(*μ*_*c,j*_*′*) such that *j*^*′*^ = arg max_*j*_ *ρ*_*c,j*_. See Web Appendix C for other hyperparameters.

### 4.4 Posterior Inference for RNA Velocity

#### 4.4.1 Gene-specific RNA Velocity

We consider two ways of estimating the RNA velocity: continuous and discrete. For a posterior MCMC draw ***θ***^(*l*)^ denoting all the parameters:

1. Compute the instant velocity 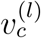 for every cell from the standard definition (Eq (1)).
2. For visualization, we also compute the predicted velocity to demonstrate cell position at one time unit ahead. Precisely, the predicted velocity 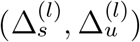 is

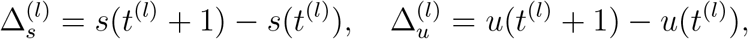

and cellular future position is 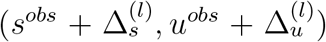. Letting ***y*** denotes the observed data, the posterior expected future position (*s*^*obs*^+ 𝔼(Δ_*s*_|***y***), *u*^*obs*^ + 𝔼(Δ_*u*_|***y***) | is approximated by

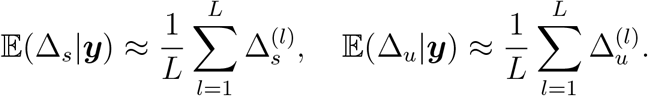

#### 4.4.2 Combined RNA Velocity

The estimation of *β*_*g*_ from the post-processing step further facilitates understanding of the dynamic information across genes, including combined RNA velocity. For each gene, we compute the future position of each cell given the same time increment (1), after applying an estimate of 1*/β*_*g*_ (posterior mean) to time-relevant quantities and *β*_*g*_ to the rates. Principal component analysis (PCA) is performed on the gene expressions combining *u* and *s*, and future total gene expression is reduced to the same lower-dimensional embeddings by applying the same principal loadings.

## 5 Simulation Study

We assess ConsensusVelo in four scenarios with large (or small) 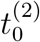 and *λ*: SS1: 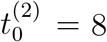, *λ* = 0.8; SS2: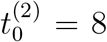, *λ* = 4; SS3: 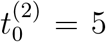, *λ* = 0.8; SS4: 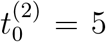, *λ* = 4, keeping all the other parameters the same. On the one hand, weak identifiability is present for large values of 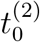 and *λ*. On the other hand, induction steady state is less likely to be reached for small values, hence affecting the empirical estimates used in the priors and initialization of the algorithm. For each setting, a single gene consisting of 200 observations with small noise (*σ*_*s*_ = *σ*_*u*_ = 1) is generated from the proposed model. For SS1, another dataset with larger noise (SS1b: *σ*_*s*_ = *σ*_*u*_ = 6) is generated for comparison. The details can be found in Web Appendix E.

For each parameter, the results are evaluated based on the posterior coverage of the 95% highest posterior density (HPD) credible intervals (CI), which we define as the proportion of cells whose true *τ*_*c*_, *t*_*c*_, *v*_*c*_, *u*_*c*_, *s*_*c*_ fall within the 95% HPD CI. Here *u*_*c*_, *s*_*c*_ denote the mean values (Eq (3)). Also, the fitted phase portrait for each posterior sample is plotted to evaluate the model fit.

### 5.1 Results from A Single Chain

The posterior coverage for *τ, t, v* is considerably sensitive to *k* when initializing the MCMC algorithm (Table 1 Method-single). Specifically, a higher correct classification rate (CCR) in initial *k* tends to yield better coverage for *τ*, and cells falling outside of the 95% HPD CI are usually in the steady states (Figure 6 in Web Appendix E), which are likely to be wrongly classified in initial *k*. Further, for all scenarios with small noise, the sampled *k* remains the same throughout sampling process, implying underestimated uncertainty.

**Table 1:** Proportion of cells whose cell-specific parameters are covered in the 95% HPD CI of the posterior samples, under four simulation settings. For simulation setting 1, comparisons are also made between a single chain and consensus velocity, and between a small (SS1a) and large noise (SS1b) in the data. Correct classification rate of initial *k* is provided in bracket.

| Simulation | Method | $\tau$ | $t$ | $v$ | $u$ | $s$ |
| --- | --- | --- | --- | --- | --- | --- |
| SS1a | single (0.835) | 0.785 | 0.835 | 0.795 | 0.93 | 0.845 |
|  | consensus velocity | **0.955** | **0.97** | **0.95** | **0.965** | **0.96** |
| SS1b | single (0.685) | 0.59 | 0.685 | 0.505 | 0.89 | 0.745 |
|  | consensus velocity | 0.805 | 0.875 | 0.83 | **0.97** | **0.925** |
| SS2 | single (0.765) | 0.735 | 0.8 | 0.76 | 0.95 | 0.815 |
| SS3 | single (0.845) | 0.78 | 0.73 | 0.765 | 0.93 | 0.83 |
| SS4 | single (0.8) | 0.75 | 0.785 | 0.755 | 0.97 | 0.755 |

Further, the accuracy of *t* in repression is highly dependent on 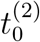 due to the relationship 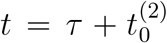, which causes a low coverage for *t* in SS3. Specifically, SS3 is the only setting where 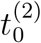 is not accurately estimated. Featured with both a small 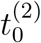 and a small *λ*, cells from SS3 are the furthest away from reaching the steady states, out of all four simulation settings, whilst our empirical estimates are calculated assuming steady states have been attained. Moreover, the mean values *u* and *s* have comparatively better posterior coverage than the other cell-wise parameters even though *τ* may be incorrect, implying weak identifiabiliy of *τ*. Generally, *u* has better coverage than *s* due to its simplest functional form, whereas *s* relies on one more parameter *γ*, which causes a low coverage in SS1b and SS4. The parameter *λ* is observed to have large uncertainty when it is large (SS2 and SS4), and 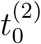 tends to have large posterior uncertainty with right-skewed distribution in all cases, both of which result from their weak identifiability.

It is worth mentioning that if the preparation step is removed and empirical estimates are used for initialization of the complete algorithm, the sampled states may be close to random in the *u − s* space with explosively large noise estimates, implying the preparation step is indispensable (Figure 7 in Web Appendix E).

### 5.2 Results from ConsensusVelo

The above results can be notably improved through the consensus velocity algorithm. Consider SS1 as an example. Utilizing posterior samples from iteration indices *i* within the range of [4901, 5000] across *W* = 100 chains yields substantial improvement (see Figure 8 in Web Appendix E for choosing *W* and *D*). The posterior coverage consistently exceeds 0.95 for all cell-wise parameters (Table 1 SS1a). This surpasses the outcomes of a single run where samples are collected for *i ∈* [15001, 25000]. Importantly, utilizing samples from later iterations does not yield any additional increase in coverage, validating the choice of *D*.

The predicted velocity (Figure 9 in Web Appendix E) is well estimated with the truth covered by the posterior samples. Further, for uncertain cells in the two corners, the sampled predicted velocity may point to opposite directions due to cells being allocated to different states in consensus velocity, showcasing the associated uncertainty as expected. On the other hand, the variability in predicted velocity is generally smaller for cells in the middle. Finally, in the presence of larger noise, the posterior coverage from a single chain can be as low as around 0.6 due to a large misclassification rate in the initialization. Applying consensus velocity with *i ∈* [4901, 5000] and *W* = 100 yields a considerable improvement to be above 0.8 (Table 1 SS1b). While the predicted velocity (Figure 9 in Web Appendix E) exhibits considerably large uncertainty, as anticipated due to increased noise in the data, the true velocity remains well within the coverage of the posterior samples.

To validate the model for inferring gene-shared latent time, we simulate multiple genes with a shared latent time and apply consensus velocity for each gene. The estimated shared latent time is close to the truth, with good coverage for small 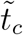, and the ordering of cells is well estimated. In addition, log(*β*_*g*_) can also be reasonably inferred (see Web Appendix E).

## 6 Application to Real Data

We demonstrate the application of ConsensusVelo in a case study on the mouse embryonic stem cells (mESCs) (Riba et al., 2022), which is enriched for proliferating cells to study the cell cycle. Genes regulated during the cell cycle are expected to show a full circular pattern in the unspliced-spliced space with dynamical information as they complete both activation and deactivation phase. We benchmark ConsensusVelo to the scVelo regarding the fitted phase portrait, and quantify the uncertainty in predicted velocities.

### 6.1 Pre-processing

The subset of genes (31) in the GOterm:cell cycle (GO:0007049) that pass the hotelling filtering (Riba et al., 2022), a process to identify of cycling genes, are selected. The data is pre-processed in scVelo to improve data quality. To align with the assumption of a truncated normal likelihood, the subset of genes with excessive zeros (*>* 1%) are further removed, leading to 26 genes for 5637 cells for model fitting.

### 6.2 Results

#### 6.2.1 Gene-specific Inference

ConsensusVelo is applied to all genes with *W* = 60 to account for the variability in cell states, and the chosen number of iterations *D* is around 5000 for most genes. When choosing the number of iterations *D*, we have also considered the absolute difference in the mean values *u* and *s*, which suggests a similar value for *D*. For each gene, 6000 MCMC samples are saved for posterior inference.

Figure 3 shows that the estimated fitted phase portrait from ConsensusVelo captures the relationship between *u* and *s* well with a complete loop. In contrast, scVelo fails to close the phase (Riba et al., 2022), especially, for gene *Top2a* that is mostly in the induction phase for scVelo. Further, assuming a zero initial condition seems incompatible with gene *Cks2*, accounting for its incomplete phase. As for gene *Nusap1* with the ideal almond shape exhibiting clear separation in the middle, scVelo infers an overestimated curvature, leading to the fit outside of the observed expression. The results from ConsensusVelo show the smallest uncertainty for *Nusap1*, aligning with its desirable shape.

The dynamics of the cells can be explored through the predicted velocity with uncertainty (Figure 3 bottom). Additionally, using a Bayesian approach allows us to evaluate the uncertainty in cell states (Figure 2 top-right), instead of a point estimate as used in scVelo. The cells in the middle of the phase portrait are more likely to belong to either of induction or repression, with the posterior probability of induction *p*_*c*_ around 0.5, whilst the classification is more certain as cells deviate from the off-diagonal line.

#### 6.2.2 Inferring the Latent Time

A linear relationship is observed between the estimated gene-specific times (Figure 4 top), even though the models for all genes are fitted independently, which suggests a potential of a shared latent time across genes for each cell. The main difference is the large values of *t* in repression which are only weakly identifiable.

Following this observation, we conduct the post-processing step to infer a gene-shared latent time 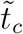 for every cell, as described in Section 4.3. Unlike frequentist estimates, the considerable amount of overlaps in the posterior densities of 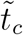 implies that it is difficult to make definite decisions on whether one cell is in an later (or earlier) stage compared to another cell (Figure 4 bottom). Instead, it would be effective to compute the posterior probability of one cell before another (Figure 4 bottom-right), offering a distinct probabilistic profile of cell orders. These probabilities can be further used as the input for downstream analysis to infer cell types (Campbell and Yau, 2019).

We compare our gene-shared latent time with transcriptional phase *θ* from DeepCycle (Riba et al., 2022), and notice that while the order of the cells appears to be similar, the two quantities are different in terms of a shift in the starting point (Figure 5 top). Further, regarding the estimated gene-shared latent time from scVelo, the starting and the end points do not seem to match. It should be noted that even though different sets of genes have been used for learning gene-shared latent time, *Nusap1* is included in all models.

The predicted velocity on the PCA space (Section 4.4.2) is shown in Figure 5 (bottom-left), where a circular pattern is observed that may be attributed to cell cycle. The direction of the velocities matches with the cell cycle phases identified in Riba et al. (2022). The same pattern is observed when examining the estimated gene-shared latent time (Figure 5 bottom-middle). Furthermore, it is also of interest to compare 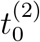 adjusted by 1*/β*_*g*_ (posterior mean) across genes (Figure 5 bottom-right). Some genes have an earlier switching time such as *Cenpe* that shows the smallest uncertainty. Gene *Cdk1* appears to have the greatest switching time, and a large uncertainty is observed in gene *Kif11* exhibiting a long right tail, indicating the possibility of switching even later than *Cdk1*.

#### 6.2.3 Posterior Predictive Checks

To examine the fit of our model, we conduct posterior predictive checks for a sample gene, *Nusap1*. A single set of the generated observations aligns with the actual data well (see Figure 14 in Web Appendix F). Further, we compute the 99% CI for each cell for *u* and *s* separately, with 1000 replicates, and 98.65% of the observed data fall inside the 99% CI in either *u* or *s* which shows appropriate estimation of uncertainty.

As for comparisons based on key statistics from the data, the empirical values of *γ* and *m*-th quantile (*m* = 5, 10, …, 95) from the replicates match the observed data well, supporting the model fit. Finally, for 1000 replicates, the posterior predictive p-values have distribution close to the uniform distribution for both states, with slightly fewer cells around 0 and 1, likely to be due to conditioning on the high probability of being in the given state, indicating a reasonable fit of our model (see Web Appendix F).

## 7 Discussion

We introduce a Bayesian hierarchical model as an alternative approach to estimate RNA velocity. This model extends the widely used method in scVelo by leveraging a time-dependent transcription rate and non-trivial initial conditions. Instead of just a point estimate, like in scVelo, our method can also output posterior distributions over all quantities of interest, allowing to evaluate their (joint) variability, including variance and credible regions, propagating the uncertainty at different levels of the considered hierarchical model. We conduct a comprehensive simulation study with various scenarios to validate our ConsensusVelo model, introducing the concept of consensus velocity to thoroughly account for the variability in the data. We present estimated phase portraits to describe the relationship between unspliced and spliced mRNA, and offer predictions of future cell positions with associated uncertainty.

Our method outperforms scVelo on the mouse embryonic stem cells by identifying phase portraits consistent with the observed gene expressions. Further, we introduce a novel post-processing step to infer gene-shared latent time, utilizing a probabilistic model for gene-specific latent time, as opposed to relying on the quantiles in scVelo. The resulting gene-shared latent time enhances our understanding of cell orders with uncertainty, supporting downstream analysis such as identifying cell populations (Campbell and Yau, 2019).

Although we consider our method using probabilistic modeling as a more rigorous approach to enhance the estimation of RNA velocity, there are still multiple practical challenges to address. First, we borrow several assumptions from scVelo, including a common starting time of the induction and independence between genes. Future work could exploit a joint model where starting time can vary across genes, and a unified gene-shared latent time is directly modeled, instead of conducting a post-processing step. However, our ConsensusVelo is more computationally efficient and still allows for inferring gene-shared latent reasonably. Secondly, pre-processing steps in scVelo, such as *k* nearest neighbours and counting of spliced and unspliced mRNAs, can significantly influence RNA velocity modeling (Gorin et al., 2022; Gao et al., 2022). It is necessary to consider the possibility of modeling raw counts (Qin et al., 2022) with minimal pre-processing, and a different data likelihood accounting for zero-inflation, e.g., negative binomial (Neelon, 2019), is worth exploring. Lastly, time-dependence can also be introduced for splicing and degradation rates beyond parametric forms. A nonparametric relationship may bring more flexibility to generalize the model, allowing it to be applied across different biological processes. However, this may introduce challenges to identifiability and interpretation. In addition, to enhance scalability, alternatives to Gibbs sampling and adaptive Metropolis-Hastings could be exploited (Fong et al., 2019). Instead of consensus velocity, Bayesian stacking (Yao et al., 2022) is another option to explore multimodal posterior distribution that weights each chain differently.

Overall, our ConsensusVelo model advances RNA velocity modeling by providing interpretable insights while accounting for uncertainty. We anticipate that this approach will be valuable to both researchers and end-users, enriching the field of RNA velocity analysis.

## Notes

### Competing Interest Statement

The authors have declared no competing interest.

## References

Barile, M., Imaz-Rosshandler, I., Inzani, I., Ghazanfar, S., Nichols, J., Marioni, J. C., Guibentif, C., and Göttgens, B. (2021). Coordinated changes in gene expression kinetics underlie both mouse and human erythroid maturation. Genome Biology, 22(1):1–22.

Battich, N., Beumer, J., de Barbanson, B., Krenning, L., Baron, C. S., Tanenbaum, M. E., Clevers, H., and van Oudenaarden, A. (2020). Sequencing metabolically labeled transcripts in single cells reveals mRNA turnover strategies. Science, 367(6482):1151–1156.

Bergen, V., Lange, M., Peidli, S., Wolf, F. A., and Theis, F. J. (2020). Generalizing RNA velocity to transient cell states through dynamical modeling. Nature Biotechnology, 38(12):1408–1414.

Bergen, V., Soldatov, R. A., Kharchenko, P. V., and Theis, F. J. (2021). RNA velocity—current challenges and future perspectives. Molecular Systems Biology, 17(8):e10282.

Campbell, K. R. and Yau, C. (2016). Order under uncertainty: robust differential expression analysis using probabilistic models for pseudotime inference. PLoS Computational Biology, 12(11):e1005212.

Campbell, K. R. and Yau, C. (2019). A descriptive marker gene approach to single-cell pseudotime inference. Bioinformatics, 35(1):28–35.

Coleman, S., Kirk, P. D., and Wallace, C. (2022). Consensus clustering for Bayesian mixture models. BMC Bioinformatics, 23(1):290.

Cui, H., Maan, H., and Wang, B. (2022). DeepVelo: Deep learning extends RNA velocity to multi-lineage systems with cell-specific kinetics. bioRxiv.

Farrell, S., Mani, M., and Goyal, S. (2022). Inferring single-cell transcriptomic dynamics with structured latent gene expression dynamics. bioRxiv.

Fong, E., Lyddon, S., and Holmes, C. (2019). Scalable nonparametric sampling from multimodal posteriors with the posterior bootstrap. In International Conference on Machine Learning, pages 1952–1962. PMLR.

Gao, M., Qiao, C., and Huang, Y. (2022). UniTVelo: temporally unified RNA velocity reinforces single-cell trajectory inference. Nature Communications, 13(1):6586.

Gayoso, A., Weiler, P., Lotfollahi, M., Klein, D., Hong, J., Streets, A. M., Theis, F. J., and Yosef, N. (2022). Deep generative modeling of transcriptional dynamics for RNA velocity analysis in single cells. bioRxiv.

Gelman, A., Meng, X.-L., and Stern, H. (1996). Posterior predictive assessment of model fitness via realized discrepancies. Statistica Sinica, pages 733–760.

Gorin, G., Fang, M., Chari, T., and Pachter, L. (2022). RNA velocity unraveled. PLOS Computational Biology, 18(9):1–55.

Griffin, J. E. and Stephens, D. A. (2013). Advances in Markov chain Monte Carlo. In Bayesian Theory and Applications. Oxford University Press, Oxford.

Gu, Y., Blaauw, D., and Welch, J. D. (2022). Bayesian inference of RNA velocity from multi-lineage single-cell data. bioRxiv.

Huang, Y. and Sanguinetti, G. (2021). Uncertainty versus variability: Bayesian methods for analysis of scRNA-seq data. Current Opinion in Systems Biology, 28:100375.

La Manno, G., Soldatov, R., Zeisel, A., Braun, E., Hochgerner, H., Petukhov, V., Lidschreiber, K., Kastriti, M. E., Lönnerberg, P., Furlan, A., Fan, J., Borm, L. E., Liu, Z., van Bruggen, D., Guo, J., He, X., Barker, R., Sundström, E., Castelo-Branco, G., Cramer, P., Adameyko, I., Linnarsson, S., and Kharchenko, P. V. (2018). RNA velocity of single cells. Nature, 560:494–498.

Lewin, A., Bochkina, N., and Richardson, S. (2007). Fully bayesian mixture model for differential gene expression: simulations and model checks. Statistical Applications in Genetics and Molecular Biology, 6(1).

Li, S., Zhang, P., Chen, W., Ye, L., Brannan, K. W., Le, N.-T., Abe, J.-i., Cooke, J. P., and Wang, G. (2023). A relay velocity model infers cell-dependent RNA velocity. Nature Biotechnology, pages 1–10.

Manuel, M., Tan, K. B., Kozic, Z., Molinek, M., Marcos, T. S., Razak, M. F. A., Dobolyi, D., Dobie, R., Henderson, B. E. P., Henderson, N. C., Chan, W. K., Daw, M. I., Mason, J. O., and Price, D. J. (2022). Pax6 limits the competence of developing cerebral cortical cells to respond to inductive intercellular signals. PLoS Biology, 20(9):1–54.

Marshall, E. C. and Spiegelhalter, D. J. (2003). Approximate cross-validatory predictive checks in disease mapping models. Statistics in Medicine, 22(10):1649–1660.

Neelon, B. (2019). Bayesian zero-inflated negative binomial regression based on pólyagamma mixtures. Bayesian Analysis, 14(3):829.

Qiao, C. and Huang, Y. (2021). Representation learning of RNA velocity reveals robust cell transitions. Proceedings of the National Academy of Sciences, 118(49):e2105859118.

Qin, Q., Bingham, E., La Manno, G., Langenau, D. M., and Pinello, L. (2022). Pyro-Velocity: Probabilistic RNA Velocity inference from single-cell data. bioRxiv, pages 2022–09.

Qiu, X., Zhang, Y., Martin-Rufino, J. D., Weng, C., Hosseinzadeh, S., Yang, D., Pogson, A. N., Hein, M. Y., Min, K. H. J., Wang, L., et al. (2022). Mapping transcriptomic vector fields of single cells. Cell, 185(4):690–711.

Riba, A., Oravecz, A., Durik, M., Jiménez, S., Alunni, V., Cerciat, M., Jung, M., Keime, C., Keyes, W. M., and Molina, N. (2022). Cell cycle gene regulation dynamics revealed by RNA velocity and deep-learning. Nature Communications, 13(1):2865.

Roberts, G. O. and Rosenthal, J. S. (2009). Examples of adaptive MCMC. Journal of Computational and Graphical Statistics, 18(2):349–367.

Tang, X., Huang, Y., Lei, J., Luo, H., and Zhu, X. (2019). The single-cell sequencing: new developments and medical applications. Cell & Bioscience, 9:1–9.

Wang, X. and Zheng, J. (2021). Velo-Predictor: an ensemble learning pipeline for RNA velocity prediction. BMC Bioinformatics, 22(10):419.

Yao, Y., Vehtari, A., and Gelman, A. (2022). Stacking for non-mixing Bayesian computations: The curse and blessing of multimodal posteriors. The Journal of Machine Learning Research, 23(1):3426–3471.

Zopf, C., Quinn, K., Zeidman, J., and Maheshri, N. (2013). Cell-cycle dependence of transcription dominates noise in gene expression. PLoS Computational Biology, 9(7):e1003161.

