## Supplementary material for "Quantifying uncertainty in RNA velocity"

**Abstract**

This supplement includes information for identifiability of the model, initialization used in the consensus velocity algorithm, empirical estimates, and details for the Gibbs sampling algorithm for posterior inference, as well as additional results for simulation study and real data analysis.

### A Fisher information and identifiability

In this section we address the issue of identifiability of time $t_c$. A simple way to do that is to compute its Fisher information which quantifies information about this parameter contained in the considered model, assuming all other parameters remain known. Therefore, in a single gene model, we only have two observations to estimate $t_c$.

Under the truncated normal error model, for $k = 1$,

$$u_c^{obs} \sim \text{Trunc-N}(u(t_c), \sigma_u^2, (0, +\infty)), \quad s_c^{obs} \sim \text{Trunc-N}(s(t_c), \sigma_s^2, (0, +\infty)).$$

Here Trunc-N $(\mu, \sigma^2, A)$ denotes the truncated normal distribution with mean $\mu$, variance $\sigma^2$ and truncation region $A$, and its density function is given by

$$f(x|\mu, \sigma^2, A) = \frac{1}{\sqrt{2\pi}\sigma F_{\mu,\sigma}(A)} \exp\left(-(x-\mu)^2/2\sigma^2\right) \mathbb{I}(x \in A),$$

where $F_{\mu,\sigma}(A)$ is the normalising factor, and $\mathbb{I}(\cdot)$ is the indicator function that takes the value of 1 if the condition insides the bracket holds true and 0 otherwise. Under the considered model,

$$\mathbb{E}(u_c^{obs}) = u(t_c) + \sigma_u \varphi(u(t_c)/\sigma_u)/\Phi(u(t_c)/\sigma_u), \quad \mathbb{E}(s_c^{obs}) = s(t_c) + \sigma_s \varphi(s(t_c)/\sigma_s)/\Phi(s(t_c)/\sigma_s).$$

The corresponding log likelihood is

$$\ell(t_c) = -0.5\sigma_u^{-2}(u_c^{obs} - u(t_c))^2 - \log\Phi(u(t_c)/\sigma_u) - 0.5\sigma_s^{-2}(s_c^{obs} - s(t_c))^2 - \log\Phi(s(t_c)/\sigma_s).$$

Differentiating wrt to $t_c$, we have

$$\begin{aligned} \ell'(t_c) &= u'(t_c)\sigma_u^{-2}(u_c^{obs} - u(t_c)) - \sigma_u^{-1}u'(t_c)\frac{\varphi(u(t_c)/\sigma_u)}{\Phi(u(t_c)/\sigma_u)} \\ &+ s'(t_c)\sigma_s^{-2}(s_c^{obs} - s(t_c)) - \sigma_s^{-1}s'(t_c)\frac{\varphi(s(t_c)/\sigma_s)}{\Phi(s(t_c)/\sigma_s)}, \end{aligned}$$

and

$$\begin{aligned}
\ell''(t_c) &= u''(t_c)\sigma_u^{-2}(u_c^{obs} - u(t_c)) - [u'(t_c)]^2\sigma_u^{-2} + s''(t_c)\sigma_s^{-2}(s_c^{obs} - s(t_c)) - [s'(t_c)]^2\sigma_s^{-2} \\
&\quad -\sigma_u^{-1}u''(t_c)\frac{\varphi(u(t_c)/\sigma_u)}{\Phi(u(t_c)/\sigma_u)} - \sigma_s^{-1}s''(t_c)\frac{\varphi(s(t_c)/\sigma_s)}{\Phi(s(t_c)/\sigma_s)} \\
&\quad -\sigma_u^{-2}[u'(t_c)]^2\frac{\varphi'(u(t_c)/\sigma_u)\Phi(u(t_c)/\sigma_u) - [\varphi(u(t_c)/\sigma_u)]^2}{[\Phi(u(t_c)/\sigma_u)]^2} \\
&\quad -\sigma_s^{-2}[s'(t_c)]^2\frac{\varphi'(s(t_c)/\sigma_s)\Phi(s(t_c)/\sigma_s) - [\varphi(s(t_c)/\sigma_s)]^2}{[\Phi(s(t_c)/\sigma_s)]^2}.
\end{aligned}$$

Using the moments of the truncated normal distribution,

$$\begin{aligned}
I(t_c) &= -\mathbb{E}(\ell''(t_c)) = [u'(t_c)]^2\sigma_u^{-2} + [s'(t_c)]^2\sigma_s^{-2} \\
&\quad -\sigma_u^{-2}[u'(t_c)]^2\varphi(u(t_c)/\sigma_u)\frac{\varphi(u(t_c)/\sigma_u) + u(t_c)\sigma_u^{-1}\Phi(u(t_c)/\sigma_u)}{[\Phi(u(t_c)/\sigma_u)]^2} \\
&\quad -\sigma_s^{-2}[s'(t_c)]^2\varphi(s(t_c)/\sigma_s)\frac{\varphi(s(t_c)/\sigma_s) + s(t_c)\sigma_s^{-1}\Phi(s(t_c)/\sigma_s)}{[\Phi(s(t_c)/\sigma_s)]^2}.
\end{aligned}$$

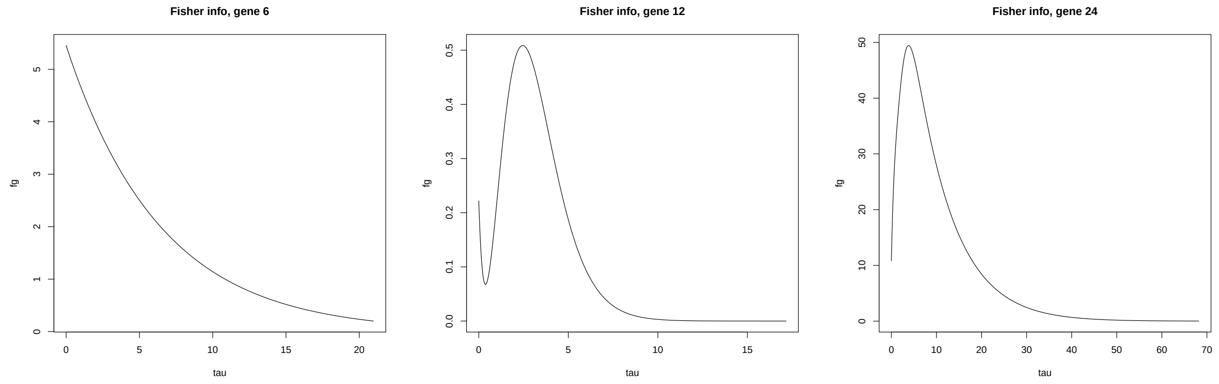

Figure 1: Fisher information as a function of $\tau$ for genes 6,12,24, for $k = 1$, $\tau \in (0, t_0^{(2)})$).

For functions $u(t)$ and $s(t)$, their first derivatives are

$$\begin{aligned}
u'(t) &= (\alpha_1^{(k)} - \beta u_0^{(k)})\exp(-\beta\tau) + \frac{\alpha_2^{(k)}}{\beta - \lambda}\left(-\beta\exp(-\beta\tau) + \lambda\exp(-\lambda\tau)\right), \\
s'(t) &= (\alpha_1^{(k)} - \gamma s_0^{(k)})\exp(-\gamma\tau) + \frac{\alpha_1^{(k)} - \beta u_0^{(k)}}{\gamma - \beta}(-\gamma\exp(-\gamma\tau) + \beta\exp(-\beta\tau)) \\
&\quad +\beta\alpha_2^{(k)}\left[\frac{-\beta\exp(-\beta\tau)}{(\beta-\lambda)(\gamma-\beta)} + \frac{\lambda\exp(-\lambda\tau)}{(\beta-\lambda)(\gamma-\lambda)} + \frac{\gamma\exp(-\gamma\tau)}{(\gamma-\beta)(\gamma-\lambda)}\right],
\end{aligned}$$

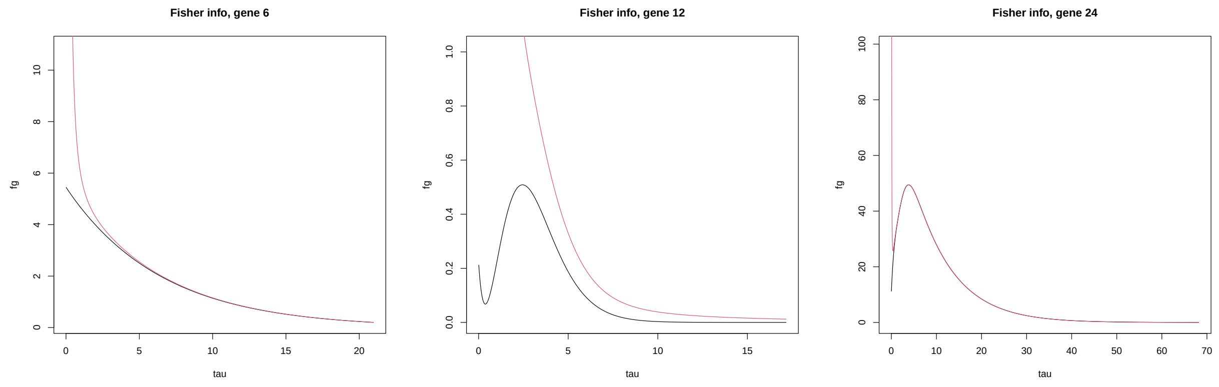

Figure 2: Fisher information as a function of $\tau$ for genes 6,12,24, for $k = 1$, $\tau \in (0, t_0^{(2)}))$, black line; with regularisation (1): red line.

where $t = \tau + t_0^{(k)}$. Note that for large $\tau$, both $u'(t)$ and $s'(t)$ tend to 0, implying that the Fisher information tends to 0, i.e. for large enough $\tau$ the model does not have information about $t_c$ - see Figure 1 for the Fisher information with plugged-in parameters from the posterior distribution for real data for genes 6, 12, 24. For genes 12 and 24, this also leads to a loss of information for small $\tau$.

Now, if we also use prior $t_c \sim \text{Gamma}(a_\tau, b_\tau)$, then the Fisher information corresponding to the log likelihood penalised by the log prior is

$$I_{pen}(t_c) = I(t_c) + \frac{a_\tau - 1}{t_c^2}. \quad (1)$$

These are plotted in Figure 2, in red. This also holds if the prior on $t_c$ is truncated to $(0, t_{max})$ with a fixed value $t_{max}$. Note that this prior distribution results in higher precision for smaller values of $\tau$.

If the prior is a gamma distribution truncated to $(0, t_{max})$ and $t_{max}$ is also estimated, e.g. by using prior $t_{max} \sim \text{Gamma}(a_0, b_0)$, then, if $a_0 \leq 1$ then the MAP of $t_{\max}$ conditionally on the remaining unknown parameters is $\max(t_c)$. Integrating out the distribution of $t_{\max}$, joint posterior distribution of $t_c$ becomes, approximately,

$$p(t_1, \dots t_n | \theta, a_\tau, b_\tau, a_0, b_0, (u_c^{obs}, s_c^{obs})) \propto (1 - F_{\Gamma(a_0,b_0)}(\max_c t_c)) \prod_c \exp(\ell(t_c, \theta)) f_{\Gamma(a_\tau,b_\tau)}(t_c),$$

thus discouraging value of $\max(t_c)$ being too large.

### B Initialization of state in consensus velocity algorithm

The primary goal of applying consensus velocity is to consider the uncertainty in the state $k$ for steady-state cells which cannot be taken into account by the Gibbs sampling algorithm. As described in the section for empirical estimates, we first use quantiles of $u$ and $s$ to determine cells that are likely to be in the steady states, denoted by $\tilde{k} = 3$ for repression steady state, and $\tilde{k} = 4$ for induction steady state. The rest cells are assigned $\tilde{k} = 1$ if $u_c^{obs} - \hat{\gamma} s_c^{obs} > 0$, and $\tilde{k} = 2$ otherwise (recall parameter is measured relative to $\beta$).

To initialize sensible $k$, we assign each cell in the steady states ($\tilde{k} > 2$) to either induction or repression phase according to cell-specific empirical probabilities. Let $p_{c,on}$ denote the empirical probability of cell $c$ belonging to induction, with $p_{c,off}$ for repression. Below we propose two ways to compute the empirical probabilities, both relying on the distance. Since the distance is sensitive to the scale, the counts $u$ and $s$ are standardized to have a mean of 0 and a standard deviation of 1, respectively. The standardization is only used for initialization, not for model fitting.

#### B.1 Distance based on the centre points in each state

Firstly, the centre point in $\tilde{k} = 1$ is calculated as the average over all data points in $\tilde{k} = 1$, denoted by $(u_{o,1}, s_{o,1})$, and the centre point for $\tilde{k} = 2$ is derived similarly, denoted by $(u_{o,2}, s_{o,2})$. For each cell $c$ in empirical steady states, the probabilities are given as follows:

$$p_{c,on} \propto \exp\left(-((u_{o,1} - u_c^{obs})^2 + (s_{o,1} - s_c^{obs})^2)\right),$$

$$p_{c,off} \propto \exp\left(-((u_{o,2} - u_c^{obs})^2 + (s_{o,2} - s_c^{obs})^2)\right).$$

An example is illustrated in Figure 3.

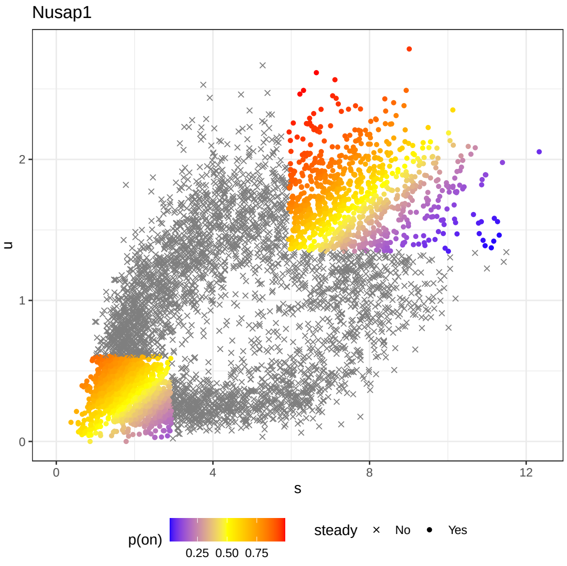

Figure 3: Empirical probabilities to initialize cells in the steady states for gene *Nusap1* based on the centre points.

#### B.2 Distance based on the closest points in each state

The centre points depend on the distribution of the cells, and hence may be biased towards one of the two steady states. Alternatively, for each steady state, we can use the closest points from $\tilde{k} = 1$ and $\tilde{k} = 2$.

For instance, let $c^*_{1,3}$ be the index of the cell in $\tilde{k} = 1$ that is the closest to the repression steady state $\tilde{k} = 3$, i.e., this point has the minimum average Euclidean distance

$$c^*_{l,m} = \arg \min_{j:\tilde{k}_j = l} \frac{1}{N_m} \sum_{i:\tilde{k}_i = m} d_{i,j},$$

where $l = 1$ or 2, $m = 3$ or 4, $N_m$ denotes the number of cells with $\tilde{k}_c = m$ and $d_{i,j}$ denotes the Euclidean distance between two cells indexed by $i$ and $j$, respectively. Given two closest points to the repression steady state ($c^*_{1,3}$ and $c^*_{2,3}$), the empirical probabilities for each cell $c$ with $\tilde{k}_c = 3$ are

$$p_{c,on} \propto \exp\left(-\frac{d^2_{c,c^*_{1,3}}}{2\sigma^2_3}\right), \quad p_{c,off} \propto \exp\left(-\frac{d^2_{c,c^*_{2,3}}}{2\sigma^2_3}\right),$$

where $\sigma_3^2$ is the variance of all Euclidean distance $d_{c,c^*_{1,3}}, d_{c,c^*_{2,3}}$, conditional on $\tilde{k}_c = 3$. The variance is applied to scale the distance because the distance based on the closest points can be very small for all cells, and therefore the change in the probability as cell moves may not be obvious. Regarding cells in the induction steady state, the probabilities depend on the identification of $c^*_{1,4}$ and $c^*_{2,4}$. An example is shown in Figure 4.

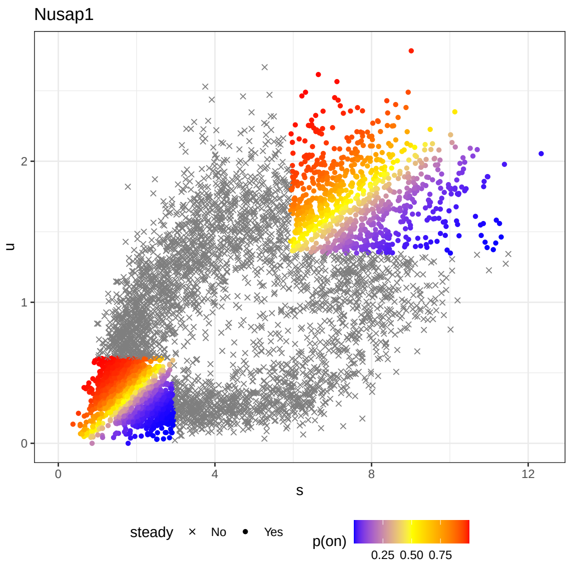

Figure 4: Empirical probabilities to initialize cells in the steady states for gene *Nusap1* based on the closest points.

### C Empirical estimates and details of prior setup

For the *ConsensusVelo* model, we estimate the initial condition for induction as:

$$u_0^- = \min(\boldsymbol{u}_{OFF}^{obs}) \cdot f_1,$$

$$u_0^+ = \max(\boldsymbol{u}_{OFF}^{obs}) \cdot f_2,$$

$$\hat{u}_0^{(1)} = \frac{1}{2}\left(u_0^- + u_0^+\right),$$

where $f_1 = 0.9$ and $f_2 = 1.1$. Note that to estimate $\tau_c$, an estimate for $u_0^{(2)}$ is also needed and is computed in a similar way to $u_0^{(1)}$ for the simulation study. For the real data, $u_0^{(2)}$ is set to $\hat{\alpha}_1^{(1)}$ and $\hat{\alpha}_1^{(1)}$ is given by the average across $\boldsymbol{u}^{obs}$ (or $\hat{\gamma}\boldsymbol{s}^{obs}$) in the estimated induction steady state, rather than the simple maximum due to the noise in the real data.

For priors of the *ConsensusVelo* model for analyzing the real data, the hyperparameters (prior means) $\mu_\alpha, \mu_\gamma, \mu_0$ are set to the empirical estimates for $\alpha, \gamma, t_0^{(2)}$, and associated standard deviations in the priors are determined by multiplying the prior mean by a factor $f = 1/6, 1/6, 1/3$, respectively. As for $\lambda$, the lower and upper abounds are $\lambda^- = 0, \lambda^+ = 5$.

In terms of hyperparameters of the priors of $\mu_{\tau,j}$ and $\sigma^2_{\tau,j}$, $\mu_j^*$ and $\eta_j$ are based on the mean and variance of empirical $\tau$ in the empirical states, with $\sigma_j^* = 2\mu_j^*$ and $\nu_j = 2\eta_j$ to allow for large variability since empirical $\tau$ is computed from the model with constant transcription rate $\alpha$, which is incorrect under the assumption of a time-dependent rate.

For the inference of gene-shared latent time from the following model:

$$\begin{aligned}
\text{Likelihood:} \quad & \bar{x}_{c,g}|\hat{k}_{c,g} = j \overset{i.i.d}{\sim} \mathrm{N}(\mu_{c,j} + \log(\beta_g), \tilde{\sigma}^2_{c,j} + \hat{\sigma}^2_{c,g,j}), \quad \hat{k}_{c,g} \overset{i.i.d}{\sim} \mathrm{Cat}(\rho_{c,1}, \rho_{c,2}), \\
\text{Priors:} \quad & \mu_{c,j} \overset{i.i.d}{\sim} \mathrm{N}(\hat{\mu}_{c,j}, \hat{\omega}^2_{c,j}), \quad \tilde{\sigma}^2_{c,j} \overset{i.i.d}{\sim} \mathrm{IG}(1, \hat{\phi}_{c,j}), \quad j = 1, 2, \\
& \rho_{c,1}, \rho_{c,2} \overset{i.i.d}{\sim} \mathrm{Dir}(\hat{\rho}_{c,1}, \hat{\rho}_{c,2}), \quad \log(\beta_g) \overset{i.i.d}{\sim} \mathrm{N}(0, \sigma_b^2),
\end{aligned} \tag{2}$$

where $\bar{x}_{c,g}$ is the posterior mean of $\log t_{c,g}$, conditional on $\hat{k}_{c,g}$, and $\hat{k}_{c,g}$ is the state with a larger posterior probability. An empirical estimate for $\log(1/\beta_g)$ $(g \neq 1)$ is obtained from the average difference between $\bar{x}_{c,1}$ and $\bar{x}_{c,g}$, where the average is taken across cells such that $\hat{k}_{c,1} = \hat{k}_{c,g}$ (recall that $\beta_1 = 1$ for identifiability). Then prior parameters $\hat{\mu}_{c,j}$ and $\hat{\phi}_{c,j}$ are given by the mean and variance of $\bar{x}_{c,g} + \log(1/\beta_g)$ with $\hat{k}_{c,g} = j$. Additionally, $\hat{\rho}_{c,j}$ is given by $G^{-1} \sum_{g=1}^{G} \hat{p}_{c,g,j}$ where $G$ is the total number of genes, and $\hat{p}_{c,g,j}$ is the posterior probability of cell $c$ belongs to state $j$ in gene $g$. The value of $\sigma_b$ is 10 for simulation study and real data analysis.

### D Posterior inference

A graphical model representation for the proposed model is shown in Figure 5.

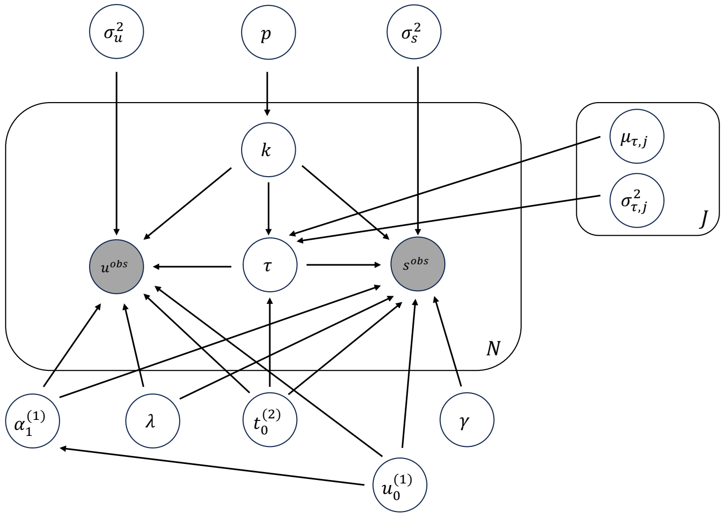

Figure 5: A directed graphical model representation of the proposed model.

Define $\boldsymbol{k} = \{k_c\}_{c=1}^C$, $\boldsymbol{\tau} = \{\tau_c\}_{c=1}^C$, $\boldsymbol{\mu}_\tau = \{\mu_{\tau,j}\}_{j=1}^2$, $\boldsymbol{\sigma}_\tau^2 = \left\{\sigma_{\tau,j}^2\right\}_{j=1}^2$. To demonstrate the dependence of the mean $u(t)$ and $s(t)$ on each parameter, we will write the mean as a function of all the necessary parameters below. The posterior distribution is

$$\pi(\boldsymbol{k}, p, \alpha_1^{(1)}, \gamma, \boldsymbol{\tau}, t_0^{(2)}, \lambda, u_0^{(1)}, \sigma_u^2, \sigma_s^2, \boldsymbol{\mu}_\tau, \boldsymbol{\sigma}_\tau^2 | \boldsymbol{u}^{obs}, \boldsymbol{s}^{obs})$$

$$\propto \prod_{c:k_c=1} \Big[\text{Trunc-N}(u_c^{obs}|u(\tau_c, \alpha_1^{(1)}, \lambda, u_0^{(1)}), \sigma_u^2, \mathbb{R}^+) \times \text{Trunc-N}(s_c^{obs}|s(\tau_c, \alpha_1^{(1)}, \lambda, u_0^{(1)}, \gamma), \sigma_s^2, \mathbb{R}^+)\Big]$$

$$\times \prod_{c:k_c=2} \Big[\text{Trunc-N}(u_c^{obs}|u(\tau_c, \alpha_1^{(1)}, \lambda, u_0^{(1)}, t_0^{(2)}), \sigma_u^2, \mathbb{R}^+) \times \text{Trunc-N}(s_c^{obs}|s(\tau_c, \alpha_1^{(1)}, \lambda, u_0^{(1)}, \gamma, t_0^{(2)}), \sigma_s^2, \mathbb{R}^+)\Big]$$

$$\times\ p^{N_1} \times (1-p)^{N_2} \times \text{Beta}(p|1,1) \times \text{Trunc-N}(\alpha_1^{(1)}|\mu_\alpha, \sigma_\alpha^2, (u_0^{(1)}, +\infty)) \times \text{Trunc-N}(\gamma|\mu_\gamma, \sigma_\gamma^2, \mathbb{R}^+)$$

$$\times \text{Unif}(\lambda|\lambda^-, \lambda^+) \times \text{Gamma}\left(t_0^{(2)}|\frac{\mu_0^2}{\sigma_0^2}, \frac{\mu_0}{\sigma_0^2}\right) \times \text{Unif}(u_0^{(1)}|u_0^-, u_0^+)$$

$$\times \prod_{c:k_c=1} \text{Trunc-Gam}\left(\tau_c|\frac{\mu_{\tau,1}^2}{\sigma_{\tau,1}^2}, \frac{\mu_{\tau,1}}{\sigma_{\tau,1}^2}, (0, t_0^{(2)})\right) \times \prod_{c:k_c=2} \text{Gamma}\left(\tau_c|\frac{\mu_{\tau,2}^2}{\sigma_{\tau,2}^2}, \frac{\mu_{\tau,2}}{\sigma_{\tau,2}^2}\right)$$

$$\times\ \text{IG}(\sigma_u^2|\xi, \xi\hat{\sigma}_u^2) \times \text{IG}(\sigma_s^2|\xi, \xi\hat{\sigma}_s^2)$$

$$\times \prod_{j=1}^{2} \left[\text{Trunc-N}(\mu_{\tau,j}|\mu_j^*, \sigma_j^{*2}, \mathbb{R}^+) \times \text{Trunc-N}(\sigma_{\tau,j}^2|\eta_j, \nu_j^2, \mathbb{R}^+)\right],$$

where $N_k = \sum_{c=1}^{C} \mathbb{I}(k_c = k)$.

#### D.1 State allocation variables $k_c$

For each cell, $k_c$ plays a role through the data likelihood, prior for $\tau_c$ and its own prior.

$$\pi(k_c = 1|u_c^{obs}, s_c^{obs}, \alpha_1^{(1)}, \gamma, \lambda, u_0^{(1)}, \tau_c, t_0^{(2)}, \sigma_u^2, \sigma_s^2, \boldsymbol{\mu}_\tau, \boldsymbol{\sigma}_\tau^2, p)$$

$$\propto \text{Trunc-N}(u_c^{obs}|u(\tau_c, \alpha_1^{(1)}, \lambda, u_0^{(1)}), \sigma_u^2, \mathbb{R}^+) \times \text{Trunc-N}(s_c^{obs}|s(\tau_c, \alpha_1^{(1)}, \lambda, u_0^{(1)}, \gamma), \sigma_s^2, \mathbb{R}^+)$$

$$\times \text{Trunc-Gam}\left(\tau_c|\frac{\mu_{\tau,1}^2}{\sigma_{\tau,1}^2}, \frac{\mu_{\tau,1}}{\sigma_{\tau,1}^2}, (0, t_0^{(2)})\right) \times p.$$

$$\pi(k_c = 2| \ldots)$$

$$\propto \text{Trunc-N}(u_c^{obs}|u(\tau_c, \alpha_1^{(1)}, \lambda, u_0^{(1)}, t_0^{(2)}), \sigma_u^2, \mathbb{R}^+) \times \text{Trunc-N}(s_c^{obs}|s(\tau_c, \alpha_1^{(1)}, \lambda, u_0^{(1)}, \gamma, t_0^{(2)}), \sigma_s^2, \mathbb{R}^+)$$

$$\times \text{Gamma}\left(\tau_c|\frac{\mu_{\tau,2}^2}{\sigma_{\tau,2}^2}, \frac{\mu_{\tau,2}}{\sigma_{\tau,2}^2}\right) \times (1-p).$$

Denote the right-hand sides as $\tilde{p}(k_c = 1)$ and $\tilde{p}(k_c = 2)$. The full conditional distribution is

$$\pi(k_c = k|\ldots) = \frac{\tilde{K}\tilde{p}(k_c = k)}{\tilde{K}\sum_{j=1}^{2}\tilde{p}(k_c = j)},$$

where $\log\left(\tilde{K}\right) = -\max_j \log\tilde{p}(k_c = j)$ is the most extreme probability that is removed to avoid computational burden when the sum in the denominator is very small. Then we just sample $k_c$ from $\{1, 2\}$ according to $\pi(k_c = k|\ldots)$. The step is repeated for every $c$.

#### D.2 Hyper-parameter $p$

The full conditional for $p$ is of a closed form:

$$\begin{aligned}\pi(p|\boldsymbol{k}) &\propto p^{N_1} \times (1-p)^{N_2} \times \text{Beta}(p|1, 1)\\ &\propto p^{N_1} \times (1-p)^{N_2},\end{aligned}$$

leading to

$$p|\boldsymbol{k} \sim \text{Beta}(N_1 + 1, N_2 + 1).$$

#### D.3 Limiting value $\alpha_1^{(1)}$ for transcriptional rate in induction

The rate $\alpha_1^{(1)}$ enters the likelihood for every cell though the mean values. For notation simplicity, we just use $u_c(\alpha_1^{(1)})$ and $s_c(\alpha_1^{(1)})$ to represent the mean values and their dependence on $\alpha_1^{(1)}$. Similar notations will be used for the remaining expositions. The full conditional for $\alpha_1^{(1)}$ is given by

$$\begin{aligned}&\pi(\alpha_1^{(1)}|\boldsymbol{u}^{obs}, \boldsymbol{s}^{obs}, \boldsymbol{\tau}, \gamma, \lambda, u_0^{(1)}, \sigma_u^2, \sigma_s^2, t_0^{(2)}, \boldsymbol{k})\\ \propto &\prod_{c=1}^{C}\left[\text{Trunc-N}(u_c^{obs}|u_c(\alpha_1^{(1)}), \sigma_u^2, \mathbb{R}^+) \times \text{Trunc-N}(s_c^{obs}|s_c(\alpha_1^{(1)}), \sigma_s^2, \mathbb{R}^+)\right]\\ &\times \text{Trunc-N}(\alpha_1^{(1)}|\mu_\alpha, \sigma_\alpha^2, (u_0^{(1)}, +\infty)),\end{aligned}$$

which is not of a standard form. Below we illustrate the steps for the AMH scheme in Algorithm 5 from Griffin and Stephens (2013).

1. Apply the following transformation to $\alpha_1^{(1)}$

$$X = g(\alpha_1^{(1)}) = \log\left(\alpha_1^{(1)} - \alpha_1^-\right) \in \mathbb{R},$$

where $\alpha_1^- = u_0^{(1)}$ is the lower bound. The Jacobian term is

$$J_x = \frac{dX}{d\alpha_1^{(1)}} = \frac{1}{\alpha_1^{(1)} - \alpha_1^-}.$$

The inverse transformation is

$$\alpha_1^{(1)} = \exp(x) + \alpha_1^-.$$

2. Algorithm 5 in Griffin and Stephens (2013) is useful to achieve a particular average acceptance probability $r$, such as 0.44 which has been shown to be optimal for a one-dimensional variable (Roberts and Rosenthal, 2009). Specifically, at iteration $n$, let $X_{old} = g(\alpha_{1,old}^{(1)})$. We propose $X_{new} \sim N(X_{old}, \zeta^n)$ where $\zeta^{(n)}$ is the adaptive variance with initial value $\zeta^{(1)} = 0.1$. It will be updated at each iteration (see step 4 below). Then $\alpha_{1,new}^{(1)}$ is obtained through inverse transformation.

3. Let $Q_n$ denote the proposal distribution at step $n$. The acceptance probability of this proposal is given by

$$\begin{aligned}\alpha(\alpha_{1,new}^{(1)}, \alpha_{1,old}^{(1)}) &= \min\left(1, \frac{\pi(\alpha_{1,new}^{(1)})Q_n(\alpha_{1,old}^{(1)}|\alpha_{1,new}^{(1)})}{\pi(\alpha_{1,old}^{(1)})Q_n(\alpha_{1,new}^{(1)}|\alpha_{1,old}^{(1)})}\right)\\ &= \min\left(1, \frac{\pi(\alpha_{1,new}^{(1)})|J_{x_{old}}|}{\pi(\alpha_{1,old}^{(1)})|J_{x_{new}}|}\right),\end{aligned}$$

where $\pi(\alpha_1^{(1)})$ is the posterior distribution. In practice, we use the log acceptance probability

$$\alpha(\alpha_{1,new}^{(1)}, \alpha_{1,old}^{(1)}) = \min\left(1, lpost_{new} - lpost_{old} + \log\left(\alpha_{1,new}^{(1)} - \alpha_1^-\right) - \log\left(\alpha_{1,old}^{(1)} - \alpha_1^-\right)\right),$$

where *lpost* denotes the posterior distribution on the log scale.

4. After making the decision to accept the proposed value or not, we now update the adaptive variance. Define

$$\omega^{(n)} = \exp\left(\log\left(\zeta^{(n)}\right) + n^{-0.7} \times \left(\alpha(\alpha_{1,new}^{(1)}, \alpha_{1,old}^{(1)}) - 0.44\right)\right),$$

then

$$\zeta^{(n+1)} = \begin{cases} \omega^{-}, & \text{if } \omega^{(n)} < \omega^{-}, \\ \omega^{(n)}, & \text{if } \omega^{(n)} \in [\omega^{-}, \omega^{+}], \\ \omega^{+}, & \text{if } \omega^{(n)} > \omega^{+}, \end{cases}$$

where $\omega^{-} = \exp(-50)$ and $\omega^{+} = \exp(50)$.

#### D.4 Degradation rate $\gamma$

Degradation rate $\gamma$ only enters the likelihood through the mean values for the spliced counts. The full conditional density is

$$\pi(\gamma|\boldsymbol{s}^{obs}, \boldsymbol{\tau}, \alpha_1^{(1)}, \lambda, u_0^{(1)}, \sigma_s^2, t_0^{(2)}, \boldsymbol{k}) \propto \prod_{c=1}^{C} \text{Trunc-N}(s_c^{obs}|s_c(\gamma), \sigma_s^2, \mathbb{R}^+) \times \text{Trunc-N}(\gamma|\mu_\gamma, \sigma_\gamma^2, \mathbb{R}^+),$$

which is not of a standard form. A log transformation is applied

$$X = \log(\gamma) \in \mathbb{R},$$

with Jacobian

$$J_x = \frac{dX}{d\gamma} = \frac{1}{\gamma},$$

and the inverse transformation is $\gamma = \exp(x)$.

Same AMH scheme is used to draw a sample for $\gamma$ as shown in Section D.3. The acceptance probability is

$$\begin{aligned}\alpha(\gamma_{new}, \gamma_{old}) &= \min\left(1, \frac{\pi(\gamma_{new})Q_n(\gamma_{old}|\gamma_{new})}{\pi(\gamma_{old})Q_n(\gamma_{new}|\gamma_{old})}\right)\\ &= \min\left(1, \frac{\pi(\gamma_{new})\gamma_{new}}{\pi(\gamma_{old})\gamma_{old}}\right)\\ &= \min\left(1, lpost_{new} - lpost_{old} + \log(\gamma_{new}) - \log(\gamma_{old})\right).\end{aligned}$$

#### D.5 Cell-specific local time $\tau_c$

The time $\tau_c$ is independent across all cells and enters the likelihood through the mean for each cell. Its full conditional density is

$$\pi(\tau_c|u_c^{obs}, s_c^{obs}, k_c, \alpha_1^{(1)}, \gamma, \lambda, u_0^{(1)}, \sigma_u^2, \sigma_s^2, t_0^{(2)}, \boldsymbol{\mu}_\tau, \boldsymbol{\sigma}_\tau^2)$$
$$\propto \text{Trunc-N}(u_c^{obs}|u_c(\tau_c), \sigma_u^2, \mathbb{R}^+) \times \text{Trunc-N}(s_c^{obs}|s_c(\tau_c), \sigma_s^2, \mathbb{R}^+) \times \text{Trunc-Gam}\left(\tau_c|\frac{\mu_{\tau,1}^2}{\sigma_{\tau,1}^2}, \frac{\mu_{\tau,1}}{\sigma_{\tau,1}^2}, (0, \tau_c^+)\right),$$

where $\tau_c^+ = t_0^{(2)}$ if $k_c = 1$ and positive infinity otherwise. The transformation used is

$$X = g(\tau_c) = -\log\left(\frac{1}{\tau_c} - \frac{1}{\tau_c^+}\right) \in \mathbb{R}.$$

The Jacobian term is

$$J_x = \frac{dX}{d\tau_c} = \frac{\tau_c^+}{\tau_c(\tau_c^+ - \tau_c)} = \frac{1}{\tau_c(1 - \tau_c/\tau_c^+)},$$

and the inverse transformation is

$$\tau_c = \frac{1}{\exp(-x) + 1/\tau_c^+} \in \left(0, \tau_c^+\right).$$

Note that for $\tau_c^+ = +\infty$, the transformation reduces to a log transformation. We still apply AMH to sample $\tau_c$. The acceptance probability is

$$\begin{aligned}\alpha(\tau_{c,new}, \tau_{c,old}) &= \min\left(1, \frac{\pi(\tau_{c,new})Q_n(\tau_{c,old}|\tau_{c,new})}{\pi(\tau_{c,old})Q_n(\tau_{c,new}|\tau_{c,old})}\right)\\ &= \min\left(1, \frac{\pi(\tau_{c,new})\tau_{c,new}(1 - \tau_{c,new}/\tau_c^+)}{\pi(\tau_{c,old})\tau_{c,old}(1 - \tau_{c,old}/\tau_c^+)}\right)\\ &= \min\Big(1, lpost_{new} - lpost_{old} + \log(\tau_{c,new}) + \log\big(1 - \tau_{c,new}/\tau_c^+\big) -\\ &\quad \log(\tau_{c,old}) - \log\big(1 - \tau_{c,old}/\tau_c^+\big)\Big).\end{aligned}$$

#### D.6 Switching time point $t_0^{(2)}$

The switching point $t_0^{(2)}$ plays a role through the means for cells in repression, and the prior for $\tau_c$ for cells in induction, together with its own prior. The full conditional distribution is

$$\begin{aligned}
&\pi(t_0^{(2)}|\boldsymbol{u}^{obs}, \boldsymbol{s}^{obs}, \boldsymbol{k}, \boldsymbol{\tau}, \alpha_1^{(1)}, \gamma, \lambda, u_0^{(1)}, \sigma_u^2, \sigma_s^2, \mu_{\tau,1}, \sigma_{\tau,1}^2) \\
\propto &\prod_{c:k_c=2} \left[\text{Trunc-N}(u_c^{obs}|u_c(t_0^{(2)}), \sigma_u^2, \mathbb{R}^+) \times \text{Trunc-N}(s_c^{obs}|s_c(t_0^{(2)}), \sigma_s^2, \mathbb{R}^+)\right] \\
\times &\prod_{c:k_c=1} \text{Trunc-Gam}\left(\tau_c|\frac{\mu_{\tau,1}^2}{\sigma_{\tau,1}^2}, \frac{\mu_{\tau,1}}{\sigma_{\tau,1}^2}, (0, t_0^{(2)})\right) \times \text{Gamma}\left(t_0^{(2)}|\frac{\mu_0^2}{\sigma_0^2}, \frac{\mu_0}{\sigma_0^2}\right).
\end{aligned}$$

Note that, if $\exists k_c = 1$, $t_0^{(2)}$ is bounded below $\max(\boldsymbol{\tau}_1)$, where $\boldsymbol{\tau}_1$ denotes all $\tau_c$ with $k_c = 1$. Otherwise it is on $\mathbb{R}^+$. Let $t_0^-$ denotes this lower bound. We apply the following transformation

$$X = g(t_0^{(2)}) = \log\left(t_0^{(2)} - t_0^-\right) \in \mathbb{R}.$$

where $t_0^- = \max(\boldsymbol{\tau}_1)$ if $\exists k_c = 1$ and 0 otherwise. The Jacobian term is

$$J_x = \frac{dX}{dt_0^{(2)}} = \frac{1}{t_0^{(2)} - t_0^-},$$

and the inverse transformation is

$$t_0^{(2)} = \exp(x) + t_0^-.$$

For $t_0^- = 0$, this is simply a log transformation. Same AMH with Algorithm 5 is applied for sampling, and the acceptance probability is

$$\begin{aligned}
\alpha(t_{0,new}^{(2)}, t_{0,old}^{(2)}) &= \min\left(1, \frac{\pi(t_{0,new}^{(2)})Q_n(t_{0,old}^{(2)}|t_{0,new}^{(2)})}{\pi(t_{0,old}^{(2)})Q_n(t_{0,new}^{(2)}|t_{0,old}^{(2)})}\right) \\
&= \min\left(1, \frac{\pi(t_{0,new}^{(2)})(t_{0,new}^{(2)} - t_0^-)}{\pi(t_{0,old}^{(2)})(t_{0,old}^{(2)} - t_0^-)}\right) \\
&= \min\left(1, lpost_{new} - lpost_{old} + \log\left(t_{0,new}^{(2)} - t_0^-\right) - \log\left(t_{0,old}^{(2)} - t_0^-\right)\right).
\end{aligned}$$

#### D.7 Newly introduced variable $\lambda$

The newly introduced variable $\lambda$ also enters the likelihood through the means for all cells. The full conditional is

$$\begin{aligned}
&\pi(\lambda|\boldsymbol{u}^{obs}, \boldsymbol{s}^{obs}, \boldsymbol{\tau}, \alpha_1^{(1)}, \gamma, u_0^{(1)}, \sigma_u^2, \sigma_s^2, t_0^{(2)}, \boldsymbol{k}) \\
\propto &\prod_{c=1}^{C} \left[\text{Trunc-N}(u_c^{obs}|u_c(\lambda), \sigma_u^2, \mathbb{R}^+) \times \text{Trunc-N}(s_c^{obs}|s_c(\lambda), \sigma_s^2, \mathbb{R}^+)\right] \\
&\times \text{Unif}(\lambda|\lambda^-, \lambda^+),
\end{aligned}$$

which is not of a standard form. Note that $\lambda$ is restricted on $(\lambda^-, \lambda^+)$, and hence we apply the following transformation:

$$X = g(\lambda) = \log\left(\frac{\lambda - \lambda^-}{\lambda^+ - \lambda}\right) \in \mathbb{R},$$

and the Jacobian term is

$$J_x = \frac{dX}{d\lambda} = \frac{\lambda^+ - \lambda^-}{(\lambda - \lambda^-)(\lambda^+ - \lambda)}.$$

The inverse transformation is

$$\lambda = \lambda^+ + \frac{\lambda^- - \lambda^+}{1 + \exp(x)}.$$

The acceptance probability is

$$\begin{aligned}
\alpha(\lambda_{new}, \lambda_{old}) &= \min\left(1, \frac{\pi(\lambda_{new})Q_n(\lambda_{old}|\lambda_{new})}{\pi(\lambda_{old})Q_n(\lambda_{new}|\lambda_{old})}\right) \\
&= \min\left(1, \frac{\pi(\lambda_{new})(\lambda_{new} - \lambda^-)(\lambda^+ - \lambda_{new})}{\pi(\lambda_{old})(\lambda_{old} - \lambda^-)(\lambda^+ - \lambda_{old})}\right) \\
&= \min\Bigg(1, lpost_{new} - lpost_{old} + \log\big(\lambda_{new} - \lambda^-\big) + \log\big(\lambda^+ - \lambda_{new}\big) - \\
&\log\big(\lambda_{old} - \lambda^-\big) - \log\big(\lambda^+ - \lambda_{old}\big)\Bigg),
\end{aligned}$$

and adaptive variance is updated as before.

#### D.8 Initial condition $u_0^{(1)}$

The initial condition $u_0^{(1)}$ enters the likelihood through the means for all cells and is also a hyperparameter in the prior for $\alpha_1^{(1)}$. The full conditional is

$$\begin{aligned}
&\pi(u_0^{(1)}|\boldsymbol{u}^{obs}, \boldsymbol{s}^{obs}, \boldsymbol{\tau}, \alpha_1^{(1)}, \gamma, \lambda, \sigma_u^2, \sigma_s^2, t_0^{(2)}, \boldsymbol{k}) \\
&\propto \prod_{c=1}^{C} \Big[\text{Trunc-N}(u_c^{obs}|u_c(u_0^{(1)}), \sigma_u^2, \mathbb{R}^+) \times \text{Trunc-N}(s_c^{obs}|s_c(u_0^{(1)}), \sigma_s^2, \mathbb{R}^+)\Big] \\
&\quad \times \text{Unif}(u_0^{(1)}|u_0^-, u_0^+) \times \text{Trunc-N}(\alpha|\mu_\alpha, \sigma_\alpha^2, (u_0^{(1)}, +\infty)).
\end{aligned}$$

The distribution has no closed form. Let $l_*^- = u_0^-$ and $u_*^+ = \min(u_0^+, \alpha_1^{(1)})$. Then $u_0^{(1)}$ is restricted on $(l_*^-, u_*^+)$, and a similar transformation to $\lambda$ is applied

$$X = g(u_0^{(1)}) = \log\left(\frac{u_0^{(1)} - l_*^-}{u_*^+ - u_0^{(1)}}\right) \in \mathbb{R},$$

and the Jacobian term is

$$J_x = \frac{dX}{du_0^{(1)}} = \frac{u_*^+ - l_*^-}{(u_0^{(1)} - l_*^-)(u_*^+ - u_0^{(1)})}.$$

The inverse transformation is

$$u_0^{(1)} = u_*^+ + \frac{l_*^- - u_*^+}{1 + \exp(x)}.$$

The acceptance probability is

$$\begin{aligned}
\alpha(u_{0,new}^{(1)}, u_{0,old}^{(1)}) &= \min\left(1, \frac{\pi(u_{0,new}^{(1)}) Q_n(u_{0,old}^{(1)}|u_{0,new}^{(1)})}{\pi(u_{0,old}^{(1)}) Q_n(u_{0,new}^{(1)}|u_{0,old}^{(1)})}\right) \\
&= \min\left(1, \frac{\pi(u_{0,new}^{(1)})(u_{0,new}^{(1)} - l_*^-)(u_*^+ - u_{0,new}^{(1)})}{\pi(u_{0,old}^{(1)})(u_{0,old}^{(1)} - l_*^-)(u_*^+ - u_{0,old}^{(1)})}\right) \\
&= \min\Bigg(1, lpost_{new} - lpost_{old} + \log\Big(u_{0,new}^{(1)} - l_*^-\Big) + \log\Big(u_*^+ - u_{0,new}^{(1)}\Big) - \\
&\log\Big(u_{0,old}^{(1)} - l_*^-\Big) - \log\Big(u_*^+ - u_{0,old}^{(1)}\Big)\Bigg),
\end{aligned}$$

and adaptive variance is updated as before.

#### D.9 Variance parameters $\sigma_u^2$ and $\sigma_s^2$

For $\sigma_u^2$, its full conditional density is given by

$$\pi(\sigma_u^2|\boldsymbol{u}^{obs}, \boldsymbol{\tau}, \alpha_1^{(1)}, \lambda, u_0^{(1)}, t_0^{(2)}, \boldsymbol{k}) \propto \prod_{c=1}^{C} \text{Trunc-N}(u_c^{obs}|u_c, \sigma_u^2, \mathbb{R}^+) \times \text{IG}(\sigma_u^2|\xi, \xi\hat{\sigma}_u^2).$$

Due to the normalizing constant in the likelihood that depends on $\sigma_u^2$, this full conditional density is still non-standard. We apply a log transformation $X = \log(\sigma_u^2)$ with Jacobian $J_x = dX/d\sigma_u^2 = 1/\sigma_u^2$. The inverse transformation is $\sigma_u^2 = \exp(x)$. Still applying the AMH scheme in Algorithm 5, we obtain the acceptance probability

$$\begin{aligned}
\alpha(\sigma_{u,new}^2, \sigma_{u,old}^2) &= \min\left(1, \frac{\pi(\sigma_{u,new}^2)Q_n(\sigma_{u,old}^2|\sigma_{u,new}^2)}{\pi(\sigma_{u,old}^2)Q_n(\sigma_{u,new}^2|\sigma_{u,old}^2)}\right) \\
&= \min\left(1, \frac{\pi(\sigma_{u,new}^2)\sigma_{u,new}^2}{\pi(\sigma_{u,old}^2)\sigma_{u,old}^2}\right) \\
&= \min\left(1, lpost_{new} - lpost_{old} + \log\big(\sigma_{u,new}^2\big) - \log\big(\sigma_{u,old}^2\big)\right).
\end{aligned}$$

As for $\sigma_s^2$, the full conditional density is

$$\pi(\sigma_s^2|\boldsymbol{s}^{obs}, \boldsymbol{\tau}, \alpha_1^{(1)}, \gamma, \lambda, u_0^{(1)}, t_0^{(2)}, \boldsymbol{k}) \propto \prod_{c=1}^{C} \text{Trunc-N}(s_c^{obs}|s_c, \sigma_s^2, \mathbb{R}^+) \times \text{IG}(\sigma_s^2|\xi, \xi\hat{\sigma}_s^2).$$

The sampling step is almost identical to $\sigma_u^2$ and hence is omitted here.

#### D.10 Hyper-parameters $\mu_{\tau,j}$ and $\sigma_{\tau,j}^2$

For each $j$ ($j = 1, 2$), we have

$$\pi(\mu_{\tau,j}|\boldsymbol{\tau}, \sigma_{\tau,j}^2, t_0^{(2)}, \boldsymbol{k}, \mu_j^*, \sigma_j^{*2}) \propto \prod_{c:k_c=j} \text{Trunc-Gam}\left(\tau_c|\frac{\mu_{\tau,1}^2}{\sigma_{\tau,1}^2}, \frac{\mu_{\tau,1}}{\sigma_{\tau,1}^2}, (0, \tau_c^+)\right) \times \text{Trunc-N}(\mu_{\tau,j}|\mu_j^*, \sigma_j^{*2}, \mathbb{R}^+),$$

where $\tau_c^+ = t_0^{(2)}$ if $j = 1$, and positive infinity otherwise. A log transformation is applied $X = \log(\mu_{\tau,j})$ with Jacobian $J_x = dX/d\mu_{\tau,j} = 1/\mu_{\tau,j}$ and inverse transformation $\mu_{\tau,j} =$

$\exp(x)$. The acceptance probability is

$$\begin{aligned}\alpha(\mu_{\tau,j}^{new}, \mu_{\tau,j}^{old}) &= \min\left(1, \frac{\pi(\mu_{\tau,j}^{new}) Q_n(\mu_{\tau,j}^{old}|\mu_{\tau,j}^{new})}{\pi(\mu_{\tau,j}^{old}) Q_n(\mu_{\tau,j}^{new}|\mu_{\tau,j}^{old})}\right) \\ &= \min\left(1, \frac{\pi(\mu_{\tau,j}^{new}) \mu_{\tau,j}^{new}}{\pi(\mu_{\tau,j}^{old}) \mu_{\tau,j}^{old}}\right) \\ &= \min\left(1, lpost_{new} - lpost_{old} + \log\big(\mu_{\tau,j}^{new}\big) - \log\big(\mu_{\tau,j}^{old}\big)\right).\end{aligned}$$

If the state is empty, the sample is drawn from the prior.

Regarding the variance, the full conditional density is given by

$$\pi(\sigma_{\tau,j}^2|\boldsymbol{\tau}, \mu_{\tau,j}, t_0^{(2)}, \boldsymbol{k}, \eta_j, \nu_j^2) \propto \prod_{c:k_c=j} \text{Trunc-Gam}\left(\tau_c|\frac{\mu_{\tau,1}^2}{\sigma_{\tau,1}^2}, \frac{\mu_{\tau,1}}{\sigma_{\tau,1}^2}, (0, \tau_c^+)\right) \times \text{Trunc-N}(\sigma_{\tau,j}^2|\eta_j, \nu_j^2, \mathbb{R}^+).$$

The sampling process is the same as $\mu_{\tau,j}$, with a log transformation. Details are omitted here.

### E Additional results for simulation study

#### E.1 A single gene

For each setting, other than $\lambda$ and $t_0^{(2)}$, the other parameters are set to the same values: $\alpha_1^{(1)} = 100, \gamma = 1.2, u_0^{(1)} = 30$, and 200 cells with $t$ equally spaced between $[0, 20]$ are generated. The variance parameters are set as $\sigma_u = \sigma_s = 1$.

Figure 6 shows that cells falling outside of the 95% HPD CI are usually in the steady states.

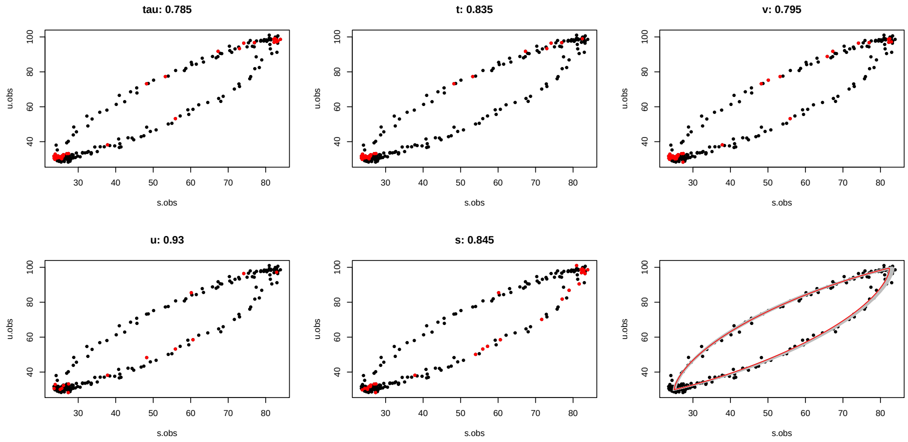

Figure 6: Cells not covered by the 95% HPD CI and uncertainty in fitted phase portrait in SS1 (small noise). The fitted phase portrait is shown in the lower-right panel (grey) with truth in red. The other panels show the cells outside of the 95% HPD CI (red), with posterior coverage in the title.

If the preparation step is removed and the empirical values of $\tau$ are used to initialize the algorithm directly, the samples for $k$ can be severely influenced and almost random. Additionally, variance of the data can get stuck into a local mode with a substantially large value (Figure 7).

Figure 8 shows the choice of $W$ and $D$ for simulation setting 1 (small noise).

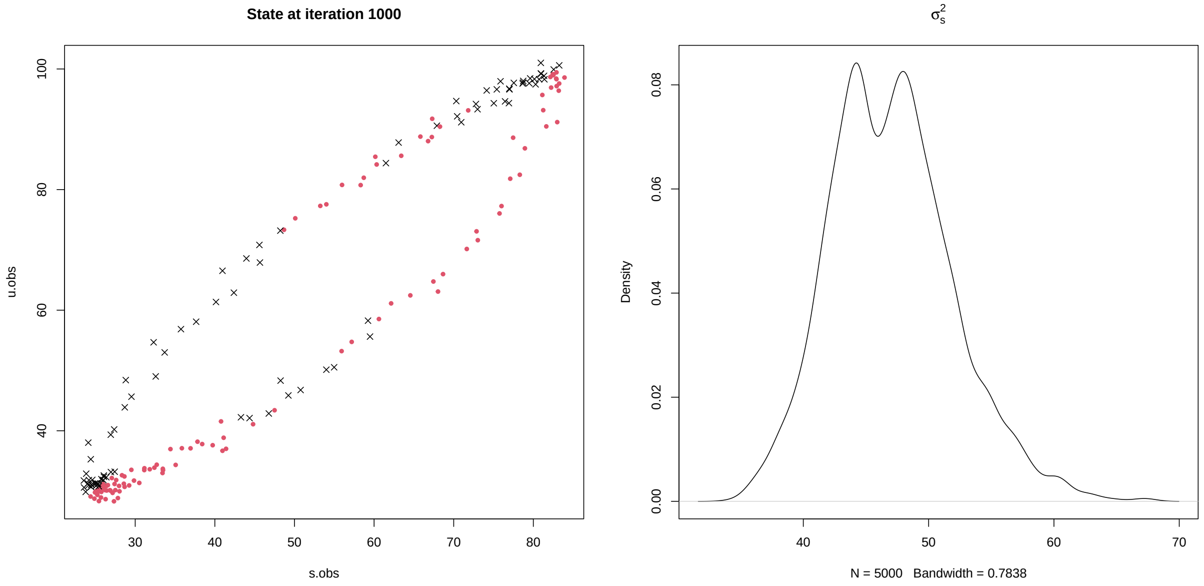

Figure 7: In SS1 (small noise), sampled $k$ at one iteration (left) and density plot for $\sigma_s^2$ (right). The true value is $\sigma_s^2 = 1$.

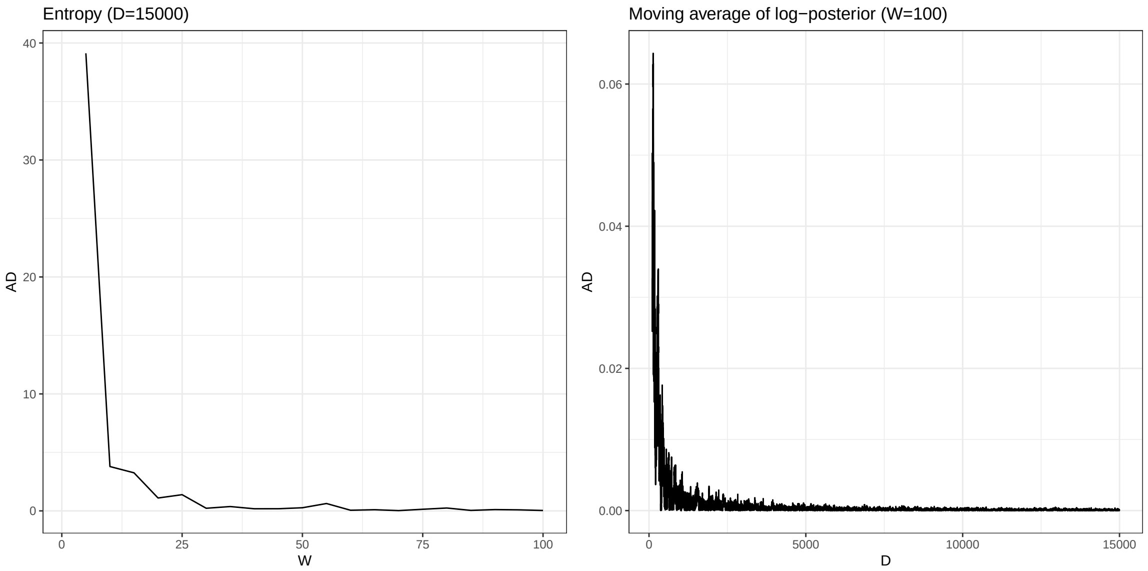

Figure 8: Choose W and D for SS1 (small noise). Left panel shows the absolute difference in entropy for $W = (5, 10, \ldots, 100)$, given $D = 15000$. Right panel shows the absolute difference in moving average of log-posterior across all 100 chains for $D = (101, 102, \ldots, 15000)$.

Figure 9 shows predicted velocity for simulation setting 1.

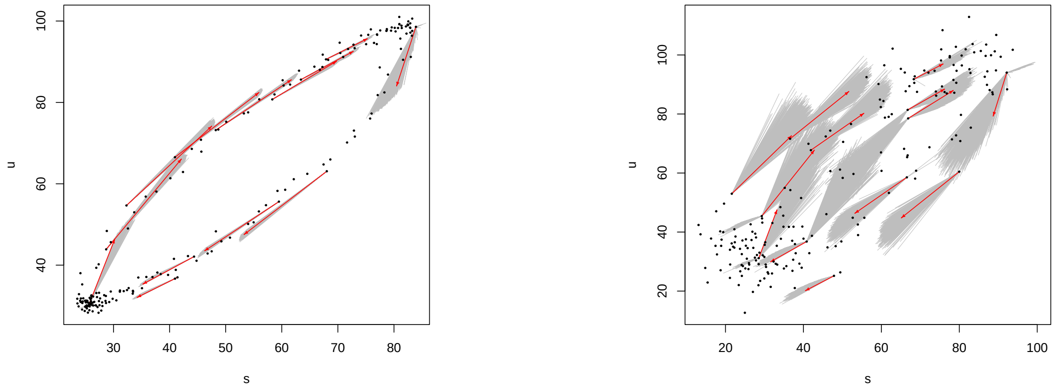

Figure 9: Predicted velocity with uncertainty for data with a small noise (left) and large noise (right) under SS1. The true velocity (red) and its uncertainty (grey) for selected cells.

#### E.2 Multiple genes

We simulate 26 genes for 400 cells with gene-shared latent time $\tilde{t}_c$ equally spaced between $[0.01, 20]$. Additionally, $1/\beta_g$ is generated from $\mathrm{Unif}(0.8, 1.2)$ and gene-wise latent time measured relative to $\beta_g$ is $t_{c,g} = \beta_g \tilde{t}_c$. The other parameters (in relative to $\beta_g$) are generated as follows:

$$\alpha_{1,g}^{(1)} \overset{i.i.d}{\sim} \text{log-N}(4.5, 0.1^2), \quad \gamma_g \overset{i.i.d}{\sim} \text{log-N}(0.1, 0.1^2),$$

$$\lambda_g \overset{i.i.d}{\sim} \mathrm{Unif}(0.6, 1.5), \quad t_{0,g}^{(2)} \overset{i.i.d}{\sim} \mathrm{N}(0.9 \times \bar{\boldsymbol{t}}_g, 0.1^2),$$

$$\sigma_{u,g}^2 \overset{i.i.d}{\sim} \text{log-N}(-0.1, 0.1^2), \quad \sigma_{s,g}^2 \overset{i.i.d}{\sim} \text{log-N}(-0.1, 0.1^2),$$

$$u_{0,g}^{(1)} \overset{i.i.d}{\sim} \text{log-N}(2, 0.1^2),$$

where $\bar{\boldsymbol{t}}_g$ is the average of gene-wise latent time across cells for gene $g$. The local time $\tau_{c,g}$ (in relative to $\beta_g$) and state $k_{c,g}$ are calculated from $t_{c,g}$ and $t_{0,g}^{(2)}$.

For each gene, consensus velocity is applied with $W = 60$ and $D = 5000$. The inferred gene-shared latent time and $\log(1/\beta_g)$ from the proposed model (Eq (2)) are shown in Figure

10. The posterior median is used as a point estimate for $\tilde{t}_c$ due to bimodal behaviour and is compared with the truth (Figure 10 left). There are points at the top-left corner, which makes sense as it is unsure whether cells from the lower-left corner of the phase portrait are at the beginning or at the end of the transcription process. The estimated shared latent time is generally close to the truth for small $\tilde{t}_c$ with small 95% CIs, and worse for large $\tilde{t}_c$ with comparatively larger CIs, which results from the difficulty in estimating $t_{0,g}^{(2)}$ (weak identifiability). Nevertheless, the ordering of the cells is well captured as suggested by the Spearman correlation between the true latent time and each posterior draw of $\tilde{t}_c$ (Figure 10 middle). As for $\beta_g$, the posterior mean is very close to the truth.

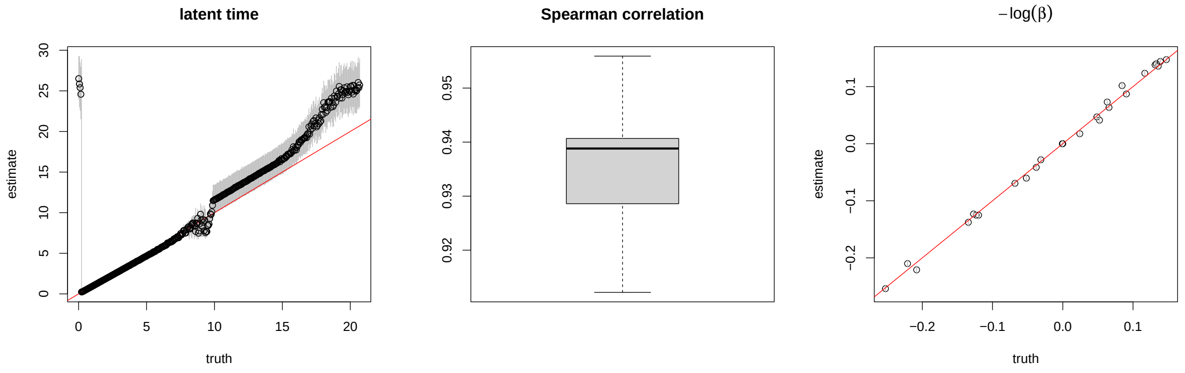

Figure 10: Estimated latent time against the truth (left), boxplot of Spearman correlation coefficients between truth and posterior samples of $\tilde{t}_c$ (middle), and posterior mean of $\log(1/\beta_g)$ against truth (right). The red line denotes $y = x$, and the grey line shows 95% CI.

### F Additional results for real data

Table 1 and Table 2 summarize posterior mean and standard deviation for parameters for 26 genes. As an example, Figure 11 showing posterior densities for one gene *Nusap1* suggests bimodal behaviours.

Table 1: Posterior means of parameters for each gene.

| | $\alpha_1^{(1)}$ | $\gamma$ | $\lambda$ | $t_0^{(2)}$ | $u_0^{(1)}$ | $\sigma_u^2$ | $\sigma_s^2$ |
|---|---|---|---|---|---|---|---|
| Aurka | 0.7592 | 0.0803 | 0.2000 | 17.0153 | 0.1317 | 0.0154 | 0.2472 |
| Bora | 0.8397 | 0.5429 | 0.2529 | 8.0934 | 0.1976 | 0.0075 | 0.0127 |
| Brca2 | 1.4114 | 0.1632 | 0.0565 | 36.1161 | 0.4203 | 0.0129 | 0.3456 |
| Ccnb1 | 0.5492 | 0.0385 | 0.1097 | 37.5365 | 0.2353 | 0.0074 | 1.2185 |
| Ccnb2 | 0.6896 | 0.0550 | 0.2238 | 14.5751 | 0.2489 | 0.0143 | 0.7726 |
| Ccnf | 2.3667 | 0.2278 | 0.0797 | 20.9771 | 0.6498 | 0.0297 | 0.2118 |
| Cdc25c | 0.6127 | 0.5201 | 0.5231 | 11.0930 | 0.1637 | 0.0083 | 0.0119 |
| Cdk1 | 0.9842 | 0.0497 | 0.0415 | 106.7518 | 0.3736 | 0.0132 | 2.2837 |
| Cenpe | 1.5535 | 0.1036 | 0.0628 | 24.1986 | 0.2966 | 0.0184 | 1.0322 |
| Cenpf | 1.3772 | 0.0543 | 0.0290 | 148.8768 | 0.2948 | 0.0171 | 4.3461 |
| Cks2 | 0.6604 | 0.0502 | 0.0953 | 31.4981 | 0.2340 | 0.0084 | 0.6589 |
| Ect2 | 3.5019 | 0.7554 | 0.5105 | 17.1694 | 1.2405 | 0.1072 | 0.2582 |
| Incenp | 0.7250 | 0.0600 | 0.0510 | 45.4799 | 0.1863 | 0.0075 | 0.5427 |
| Kif11 | 1.9202 | 0.2101 | 0.0630 | 29.0156 | 0.3851 | 0.0225 | 0.7474 |
| Kif20a | 0.4602 | 0.0548 | 0.0752 | 23.3466 | 0.1411 | 0.0054 | 0.4138 |
| Kif23 | 0.8287 | 0.2691 | 0.2247 | 10.0305 | 0.1190 | 0.0147 | 0.0371 |

| | | | | | | | |
|---|---|---|---|---|---|---|---|
| Ndc80 | 0.9986 | 0.2198 | 0.0827 | 22.1654 | 0.2489 | 0.0119 | 0.0786 |
| Nusap1 | 1.8116 | 0.1789 | 0.4345 | 12.1107 | 0.2920 | 0.0489 | 0.5279 |
| Prc1 | 1.6633 | 0.1359 | 0.0712 | 54.4076 | 0.2557 | 0.0310 | 0.4961 |
| Pttg1 | 0.5459 | 0.0745 | 0.0373 | 56.2256 | 0.0909 | 0.0055 | 0.1031 |
| Racgap1 | 0.8077 | 0.1700 | 0.2552 | 12.9787 | 0.2934 | 0.0157 | 0.1331 |
| Spdl1 | 0.8158 | 0.1720 | 0.0720 | 15.6099 | 0.1987 | 0.0064 | 0.0618 |
| Tipin | 3.2539 | 0.1435 | 0.0669 | 15.4132 | 1.3374 | 0.0464 | 1.0044 |
| Top2a | 5.0915 | 0.1387 | 0.0617 | 68.1860 | 1.4842 | 0.1590 | 12.7740 |
| Tpx2 | 1.9527 | 0.1215 | 0.3687 | 13.9866 | 0.5454 | 0.0507 | 1.3875 |
| Ttk | 1.5617 | 0.3766 | 0.1269 | 9.7021 | 0.3744 | 0.0206 | 0.0521 |

Table 2: Posterior standard deviations of parameters for each gene.

| | $\alpha_1^{(1)}$ | $\gamma$ | $\lambda$ | $t_0^{(2)}$ | $u_0^{(1)}$ | $\sigma_u^2$ | $\sigma_s^2$ |
|---|---|---|---|---|---|---|---|
| Aurka | 0.0088 | 0.0022 | 0.0123 | 0.3846 | 0.0067 | 0.0005 | 0.0223 |
| Bora | 0.0503 | 0.0079 | 0.0342 | 2.8856 | 0.0152 | 0.0005 | 0.0015 |
| Brca2 | 0.0579 | 0.0018 | 0.0057 | 4.8270 | 0.0234 | 0.0007 | 0.0241 |
| Ccnb1 | 0.0083 | 0.0006 | 0.0078 | 1.8098 | 0.0031 | 0.0003 | 0.0485 |
| Ccnb2 | 0.0295 | 0.0011 | 0.0491 | 0.9164 | 0.0045 | 0.0006 | 0.0553 |
| Ccnf | 0.0783 | 0.0028 | 0.0048 | 3.0872 | 0.0252 | 0.0015 | 0.0383 |
| Cdc25c | 0.0289 | 0.0083 | 0.1015 | 4.4769 | 0.0129 | 0.0004 | 0.0012 |
| Cdk1 | 0.0123 | 0.0005 | 0.0043 | 9.6007 | 0.0212 | 0.0010 | 0.1462 |
| Cenpe | 0.0585 | 0.0010 | 0.0051 | 1.5003 | 0.0138 | 0.0012 | 0.0862 |
| Cenpf | 0.0358 | 0.0007 | 0.0036 | 15.9777 | 0.0321 | 0.0011 | 0.7504 |

| | | | | | | | |
|---|---|---|---|---|---|---|---|
| Cks2 | 0.0185 | 0.0005 | 0.0082 | 2.3332 | 0.0040 | 0.0004 | 0.0458 |
| Ect2 | 0.1866 | 0.0081 | 0.1525 | 4.3022 | 0.0897 | 0.0093 | 0.0229 |
| Incenp | 0.0159 | 0.0009 | 0.0050 | 4.6533 | 0.0073 | 0.0003 | 0.0632 |
| Kif11 | 0.1443 | 0.0036 | 0.0068 | 3.1080 | 0.0304 | 0.0037 | 0.0590 |
| Kif20a | 0.0186 | 0.0009 | 0.0105 | 2.2474 | 0.0047 | 0.0002 | 0.0315 |
| Kif23 | 0.0494 | 0.0054 | 0.0354 | 0.4960 | 0.0113 | 0.0006 | 0.0074 |
| Ndc80 | 0.0468 | 0.0029 | 0.0084 | 3.7740 | 0.0174 | 0.0005 | 0.0081 |
| Nusap1 | 0.0332 | 0.0023 | 0.0403 | 0.5561 | 0.0115 | 0.0030 | 0.0254 |
| Prc1 | 0.1087 | 0.0053 | 0.0088 | 21.1962 | 0.0126 | 0.0023 | 0.1369 |
| Pttg1 | 0.0227 | 0.0012 | 0.0031 | 7.9332 | 0.0058 | 0.0002 | 0.0289 |
| Racgap1 | 0.0187 | 0.0018 | 0.0354 | 0.4287 | 0.0082 | 0.0006 | 0.0066 |
| Spdl1 | 0.0342 | 0.0019 | 0.0067 | 0.9603 | 0.0115 | 0.0003 | 0.0069 |
| Tipin | 0.1199 | 0.0008 | 0.0064 | 0.3741 | 0.0153 | 0.0015 | 0.0574 |
| Top2a | 0.1526 | 0.0030 | 0.0037 | 10.2239 | 0.0315 | 0.0205 | 1.4575 |
| Tpx2 | 0.0238 | 0.0012 | 0.0416 | 0.3692 | 0.0131 | 0.0018 | 0.1116 |
| Ttk | 0.1092 | 0.0058 | 0.0119 | 2.5982 | 0.0178 | 0.0013 | 0.0046 |

#### F.1 Gene-specific latent time

The posterior samples of gene-specific latent time can be obtained from $\tau, t_0^{(2)}$ and $k$. Due to the bimodal behaviour in $\tau$ and $k$, the time $t_{c,g}$ for cell $c$ in gene $g$ is also bimodal, and hence the posterior mode (or median) is more suited to provide a point estimate, rather than posterior mean. With latent time, it is possible to understand the order of the cells within each gene in the unspliced-spliced space (Figure 12), as well as the temporal change of standardized total gene expressions $u + s$ (Figure 13).

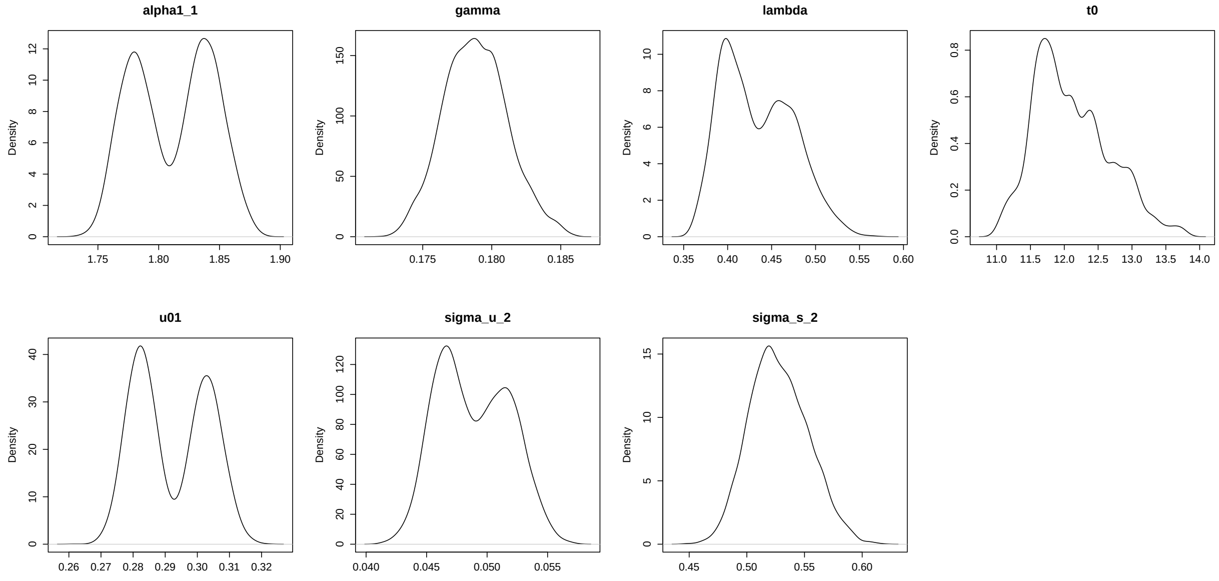

Figure 11: Posterior densities of parameters for gene *Nusap1*.

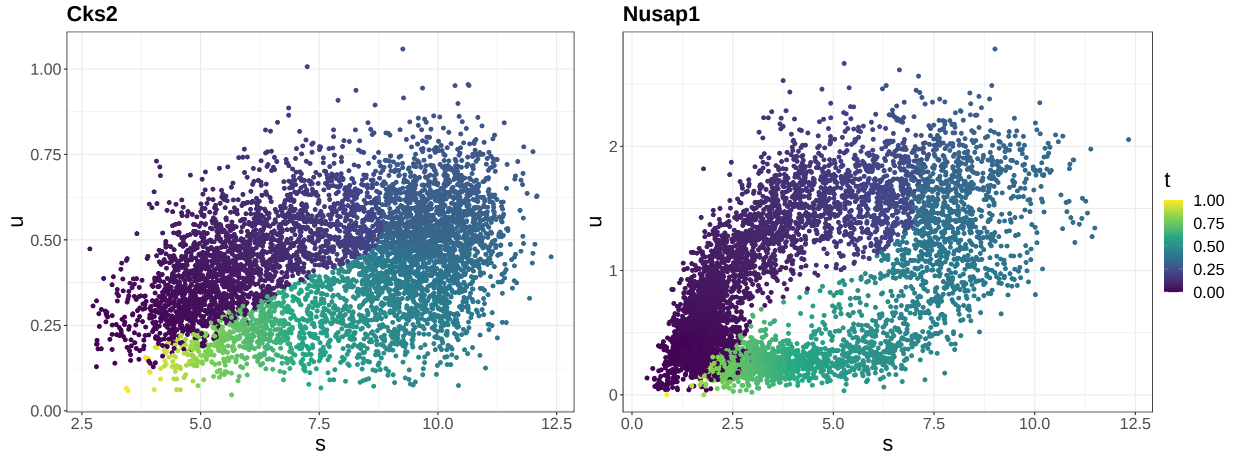

Figure 12: The estimated gene-specific latent time for example genes. Each panel shows the phase portrait with cells colored by latent time (normalized to be between 0 and 1).

#### F.2 Posterior predictive checks

For a single set of generated data, Figure 14 shows that the replicated data aligns with the actual data well. For 1000 replicates, 98.65% of the observed data fall inside the 99% CI in either $u$ or $s$ (Figure 14 bottom) implying appropriate estimation of uncertainty.

Figure 15 compares empirical $\gamma$ and quantiles from the replicates with the observed

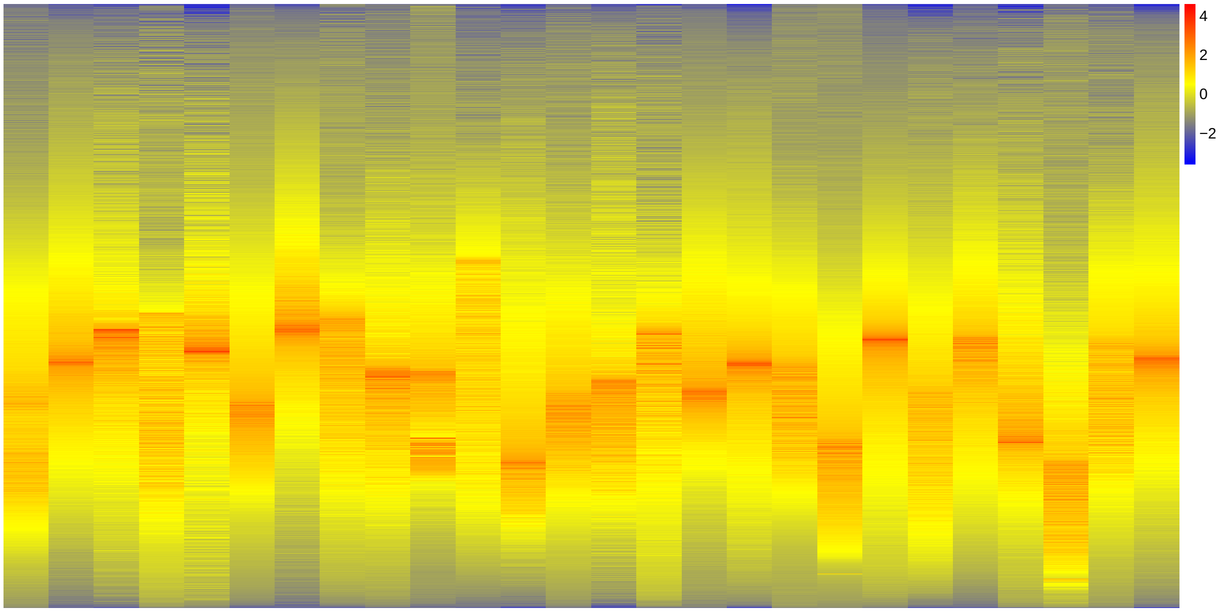

Figure 13: The total gene expressions along gene-specific latent time. Each column shows standardized gene expressions for one gene, in the order of increasing latent time from top to bottom.

data, and shows the distribution of posterior predictive p-values.

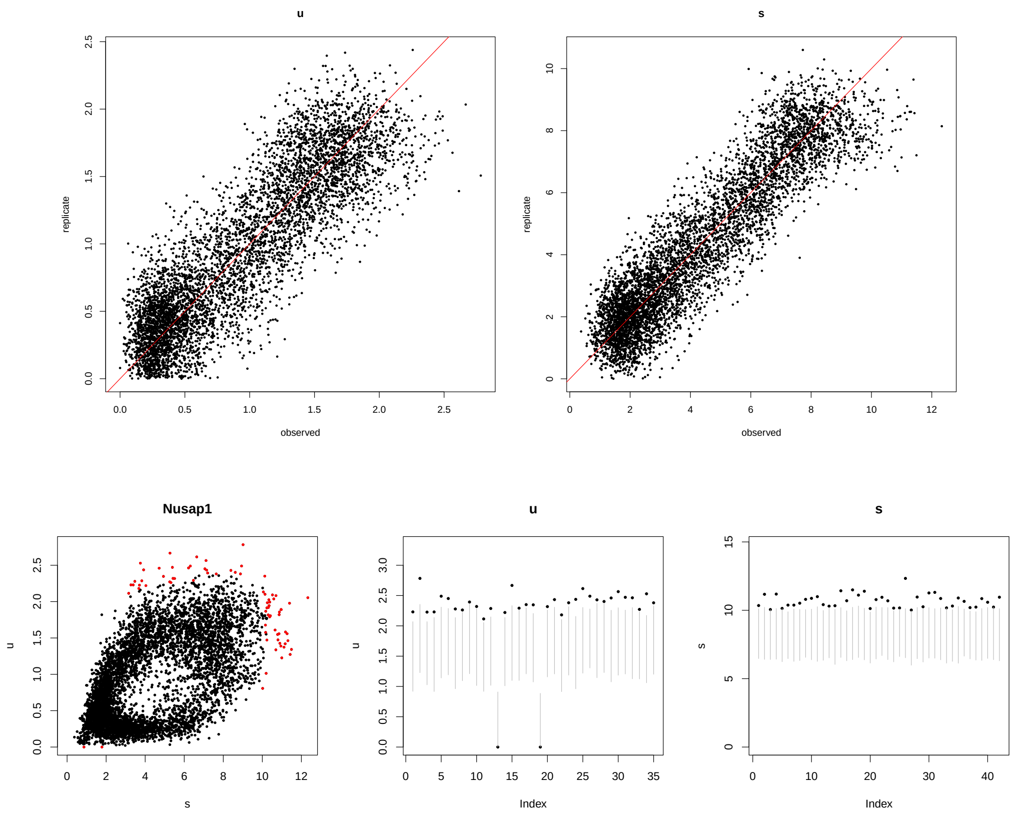

Figure 14: Top: Compare $u$ (left) and $s$ (right) in one replicated data with the observed data for gene *Nusap1*. The red line denotes $y = x$. Bottom: Cells that fall outside of the 99% CI for gene *Nusap1*. The bottom-left panel shows that only cells with extreme values (red) fall outside of the 99% CI. The middle and right panels plot the 99% CI (grey) for these cells and their observed values (black).

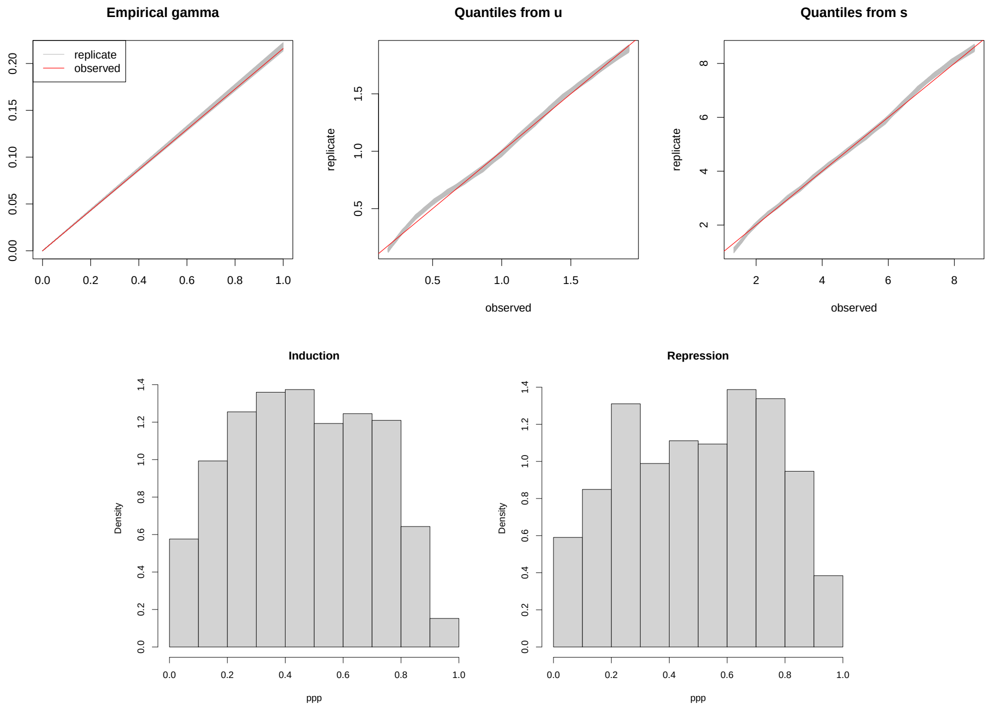

Figure 15: Top: Posterior predictive checks based on empirical $\gamma$ and quantiles. Top-left: Empirical $\gamma$ shown as the slope of the line, with the replicates in grey and observed data in red. Top-middle and top-right: Percentiles from the replicates and the actual data based on $u$ and $s$, respectively. The red line denotes $y = x$. Bottom: Posterior predictive p-values for cells with $> 0.9$ posterior probability in the given state (Bottom-left: induction, bottom-right: repression).

### References

Griffin, J. E. and Stephens, D. A. (2013). Advances in Markov chain Monte Carlo. In *Bayesian Theory and Applications*. Oxford University Press, Oxford.

Roberts, G. O. and Rosenthal, J. S. (2009). Examples of adaptive MCMC. *Journal of Computational and Graphical Statistics*, 18(2):349–367.
